# Cross-species analysis identifies APOBEC3-associated spliceosomal snRNA hypermutability

**DOI:** 10.64898/2026.10.12.758683

**Authors:** Jing Xue, Zongjin Jiang, Jilin Zhang, Cai Li

## Abstract

Spliceosomal small nuclear RNA (snRNA) genes show unusually high germline mutation rates, with R-loop formation previously implicated in this mutability. Our cross-species rare-variant and *de novo* mutation analyses show that hypermutability is concentrated in RNA polymerase II-transcribed spliceosomal snRNA genes, with strong enrichment in primates but a much weaker excess in rodents and chicken, consistent with lineage differences in APOBEC3 repertoires. Human Pol II-transcribed spliceosomal snRNA genes show mutational strand asymmetry, APOBEC3-associated mutational signatures and strong APOBEC3B binding signals. R-loops are also frequent at these loci, including in mouse genes with much weaker hypermutability, indicating that R-loop formation alone is insufficient to explain the phenotype. Together, cross-species comparisons support lineage-specific APOBEC3 activity, rather than R-loop formation alone, as a major contributor to spliceosomal snRNA gene hypermutability.

## Introduction

Spliceosomal uridine-rich small nuclear RNA (snRNA) genes, here termed *spliceRNUs*, encode core RNA components of the spliceosome that mediate splice-site recognition and intron removal^1,2^. In recent years, *spliceRNUs* have emerged as an important class of disease genes, causing predominantly developmental disorders, particularly neurodevelopmental disorders, as well as retinal disease^3–10^. Notably, many *spliceRNUs* show unusually high germline mutation rates^11,12^. However, the distribution of this hypermutability across *spliceRNU* classes and the underlying mutational processes remain unclear.

A previous mutation-rate study discussed *RNU* hypermutability in the context of RNA polymerase (Pol) III transcription^11^. However, *spliceRNUs* are transcribed by two different RNA polymerases: U1, U2, U4, U5, U11, U12 and U4atac genes are transcribed by RNA Pol II, whereas U6 and U6atac genes are transcribed by RNA Pol III^13–15^. It therefore remains unclear whether elevated mutability is shared across polymerase classes or is specific to Pol II-transcribed *spliceRNUs* (*Pol II spliceRNUs* for short). R-loops provide one plausible source of mutagenic substrate at *spliceRNUs*. These RNA-DNA hybrids expose the displaced non-template DNA strand as single-stranded DNA (ssDNA), and a recent study reported an excess *de novo* mutations (DNMs) in R-loop-forming genomic regions, with enrichment in snRNA genes^4^. This association implicates R-loop-forming loci in human germline mutagenesis but does not reveal the underlying mutagenic effector.

APOBEC3 cytidine deaminases are strong candidates because they can deaminate cytosines in host ssDNA while functioning in antiviral and anti-retroelement defence^16,17^. Human germline variation and DNM data exhibit patterns consistent with APOBEC3 activity^18,19^, and APOBEC3s can act on the displaced DNA strand of R-loops^17,20^. Thus, R-loops may create susceptible ssDNA, whereas APOBEC3 access and lesion processing may determine whether that substrate becomes mutagenic. Whether APOBEC3 activity contributes to *spliceRNU* hypermutability remains unresolved. Evolutionary comparisons provide a unique perspective on this phenomenon. APOBEC3 repertoires have undergone substantial lineage-specific expansion and diversification across mammals^21^, with corresponding differences in enzyme composition, activity, and access to cellular DNA^22,23^. If hypermutability of *spliceRNUs* depends on APOBEC3 activity rather than on R-loop formation alone, its magnitude need not be conserved across species.

Here, we integrate germline rare-variant and DNM data with cross-species comparisons and functional genomic maps to investigate *spliceRNU* hypermutability (**Fig. 1a**). We show that Pol II-transcribed *spliceRNUs* exhibit markedly elevated germline mutagenesis in human and several other primates, whereas the excess is much weaker in rodents and chicken. Mutational and functional genomic analyses further support APOBEC3-mediated deamination as a major contributor, with R-loops likely providing ssDNA substrates but insufficient on their own to explain the hypermutability.

**Fig. 1:**
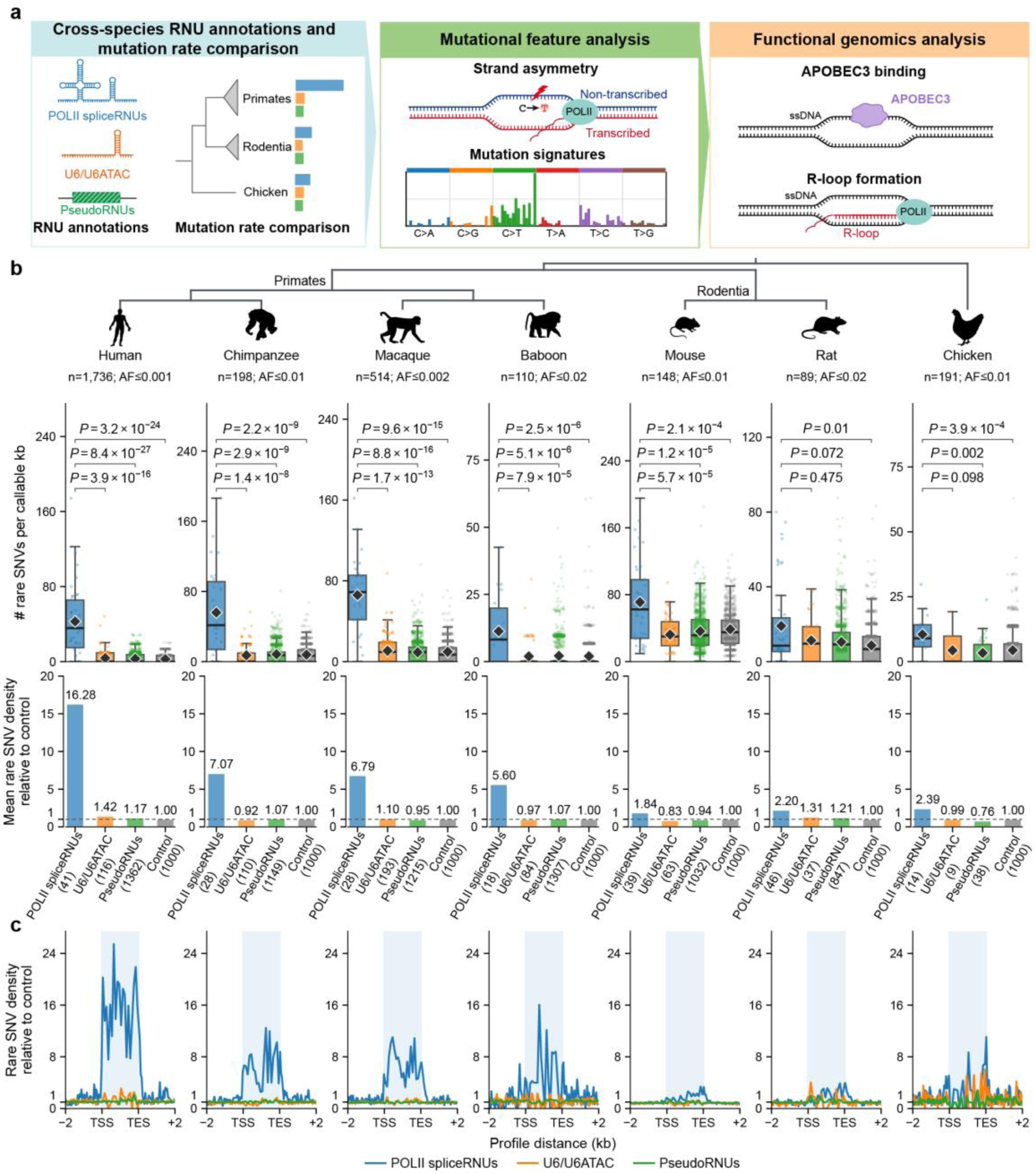
Rare SNV density and control-normalized enrichment profiles across *RNU* classes in representative species. **a**, Schematic of the study design and analytical workflow. **b**, Upper gene-level rare SNV density boxplots for human, chimpanzee, macaque, baboon, mouse, rat and chicken. Phylogenetic relationships, silhouettes, species names and rare-variant AF thresholds adjusted for species-specific sample sizes are shown above. Centre lines indicate medians; boxes, interquartile ranges; whiskers, the most extreme observations within 1.5 × the interquartile range; points, individual genes or loci; and diamonds indicate arithmetic means. Brackets compare *Pol II spliceRNUs* with each comparator and report *P* values from two-sided Mann-Whitney *U* tests. Lower barplots show mean locus density divided by the corresponding mean control density; dashed lines indicate a ratio of 1. Mean density for each class is divided by the mean density of the same species. Control loci, defined as 1,000 randomly selected, non-overlapping 150-bp intergenic regions; control equals one. **c**, Rare SNV profiles normalized to same-species control density. Gene bodies are rescaled to 1 kb (20 proportional bins), with 2-kb upstream and downstream flanks (40 bins of 50 bp each). Shading denotes the gene body. TSS, transcription start site; TES, transcription end site.

## Results

### Pol II-transcribed *spliceRNUs* exhibit hypermutability in human genomes

To identify actively transcribed spliceosomal *RNU* loci associated with Pol II, we integrated two criteria: POLR2A occupancy and candidate cis-regulatory element (cCRE) annotations^6,13^ (**Supplementary Fig. 1a**; **Supplementary Table 1**; Methods). Their intersection defined 44 high-confidence *Pol II spliceRNUs*, of which 41 with ≥50 effective callable bases were retained for downstream analysis. After applying the same callability filter, the comparison sets comprised 116 putatively functional U6/U6ATAC genes, 1,362 *RNU* pseudogenes and 1,000 autosomal 150-bp intergenic control regions in human genome (**Supplementary Fig. 1b**; **Supplementary Table 2**). We leveraged the density of rare single-nucleotide variants (SNVs) as a proxy for relative germline mutation rates^24^. At highly mutable loci, recurrent mutations in large samples can weaken the correspondence between rare variants and *de novo* mutation rate^25,26^. We therefore used the moderately sized gnomAD v4 Ashkenazi Jewish (ASJ) population as the primary dataset (1,736 individuals; AF ≤ 1 × 10^−3^)^27^.

*Pol II spliceRNUs* showed 11.49-, 13.93- and 16.28-fold higher rare SNV densities than U6/U6ATAC genes, *RNU* pseudogenes and intergenic controls, respectively (two-sided Mann-Whitney *U* tests; *P* < 1 × 10^−15^ for all tests, **Fig. 1b**). By contrast, none of the pairwise comparisons among U6/U6ATAC genes, *RNU* pseudogenes and intergenic controls was significant. The same pattern was reproduced in three additional gnomAD groups—East Asian (EAS; *n* = 2,598), South Asian (SAS; *n* = 2,417) and Finnish (FIN; *n* = 5,316)—and in an independently assembled, a fixed 2,000-individual randomly selected subset of the gnomAD v3.1.2 Human Genome Diversity Project and 1000 Genomes Project (HGDP+1KG) cohort (random2000; *n* = 2,000; **Supplementary Fig. 2**). Germline DNM analysis in 9,652 probands from Pálsson et al.^28^ independently showed a 25.63-fold higher aggregate mutation rate at *Pol II spliceRNUs* than at intergenic controls (**Supplementary Fig. 3**). Together, the rare variant and DNM analyses support markedly elevated germline mutability at human *Pol II spliceRNUs*.

### Stronger hypermutability of *Pol II spliceRNUs* in primates than in rodents and chicken

To place the human pattern in an evolutionary context, we analyzed the corresponding *RNU* classes in five other mammalian species (chimpanzee, macaque, baboon, mouse and rat), with chicken as an outgroup. In the three non-human primates, we annotated and classified *RNU* loci using regulatory chromatin annotations and synteny with human *RNU* loci (**Supplementary Fig. 1 and 4**; **Supplementary Table 1**; Methods). We identified 28, 28 and 18 *Pol II spliceRNUs* in chimpanzee, macaque and baboon, respectively, whereas the numbers of U6/U6ATAC genes and *RNU* pseudogenes were broadly comparable to those in humans (**Supplementary Table 3-5**). All three non-human primates showed significantly higher rare SNV densities at *Pol II spliceRNUs* than at U6/U6ATAC genes, *RNU* pseudogenes and intergenic controls (5.23–7.72-fold; two-sided Mann–Whitney *U* tests; *P* < 1 × 10^-4^; **Fig. 1b**), recapitulating the human pattern. Thus, the pronounced excess was shared across the four sampled primates, encompassing both apes and Old World monkeys.

For rodents and chicken, we reannotated *RNU* loci using regulatory chromatin annotations and retained loci with at least 50 effective callable bases (**Supplementary Fig. 5-6**; **Supplementary Table 1**; Methods). The final sets comprised 39, 46 and 14 *Pol II spliceRNUs* in mouse, rat and chicken, respectively (**Supplementary Table 6-8**). In mouse and rat, rare SNV densities at *Pol II spliceRNUs* were 1.69–2.20-fold higher than those at the other *RNU* classes and intergenic controls (**Fig. 1b**), representing substantially weaker enrichment than in primates. In mouse, all comparisons were significant (*P* < 1 × 10^-3^), whereas in rat, enrichment was significant relative to intergenic controls (*P* = 0.010), but not relative to U6/U6ATAC genes or *RNU* pseudogenes (*P* = 0.475 and 0.072, respectively).

Chicken also showed weaker enrichment than primates, with rare SNV densities at *Pol II spliceRNUs* elevated 2.39–3.14-fold relative to the other *RNU* classes and intergenic controls. Enrichment was significant relative to *RNU* pseudogenes and intergenic controls (*P* < 0.05), but not relative to U6/U6ATAC genes (*P* = 0.098; **Fig. 1b**). Metagene profiles showed that the excess rare SNV density was concentrated within annotated gene bodies (**Fig. 1c**). Moreover, the stronger human enrichment persisted after downsampling to 100-500 individuals (**Supplementary Fig. 7**), suggesting that sample size alone does not account for the difference. Recent exponential population growth in humans may further amplify rare-variant enrichment relative to other primates^29^. Together, these analyses show that rare SNV enrichment at *Pol II spliceRNUs* is more pronounced in the surveyed primates than in rodents or chicken.

### Mutational features support a contribution of APOBEC3 to the hypermutability of *Pol II spliceRNUs*

The stronger mutational excess at *Pol II spliceRNUs* in primates than in rodents suggests a lineage-dependent component. Previous studies reported larger APOBEC3 gene repertoires in primates than in rodents and chicken (**Fig. 2a**; **Supplementary Table 9**)^21,30–32^. Given the transcription-associated ssDNA at these actively expressed loci, APOBEC3 is a particularly plausible candidate: each primate species in our analysis encodes 6-9 canonical APOBEC3 genes, whereas rodents encode a single *Apobec3*^21,30^, with lower catalytic activity and predominantly cytoplasmic localization, potentially limiting its mutagenic activity on host DNA. Consistent with this difference, endogenous APOBEC3 mutational signatures are prominent in human cancers but have not been reported in murine cancer models^33,34^. We therefore tested for mutational features associated with off-target APOBEC3 activity, including transcriptional strand asymmetry, trinucleotide substitution spectra and rare SNV density at TCW motifs^35,36^.

**Fig. 2:**
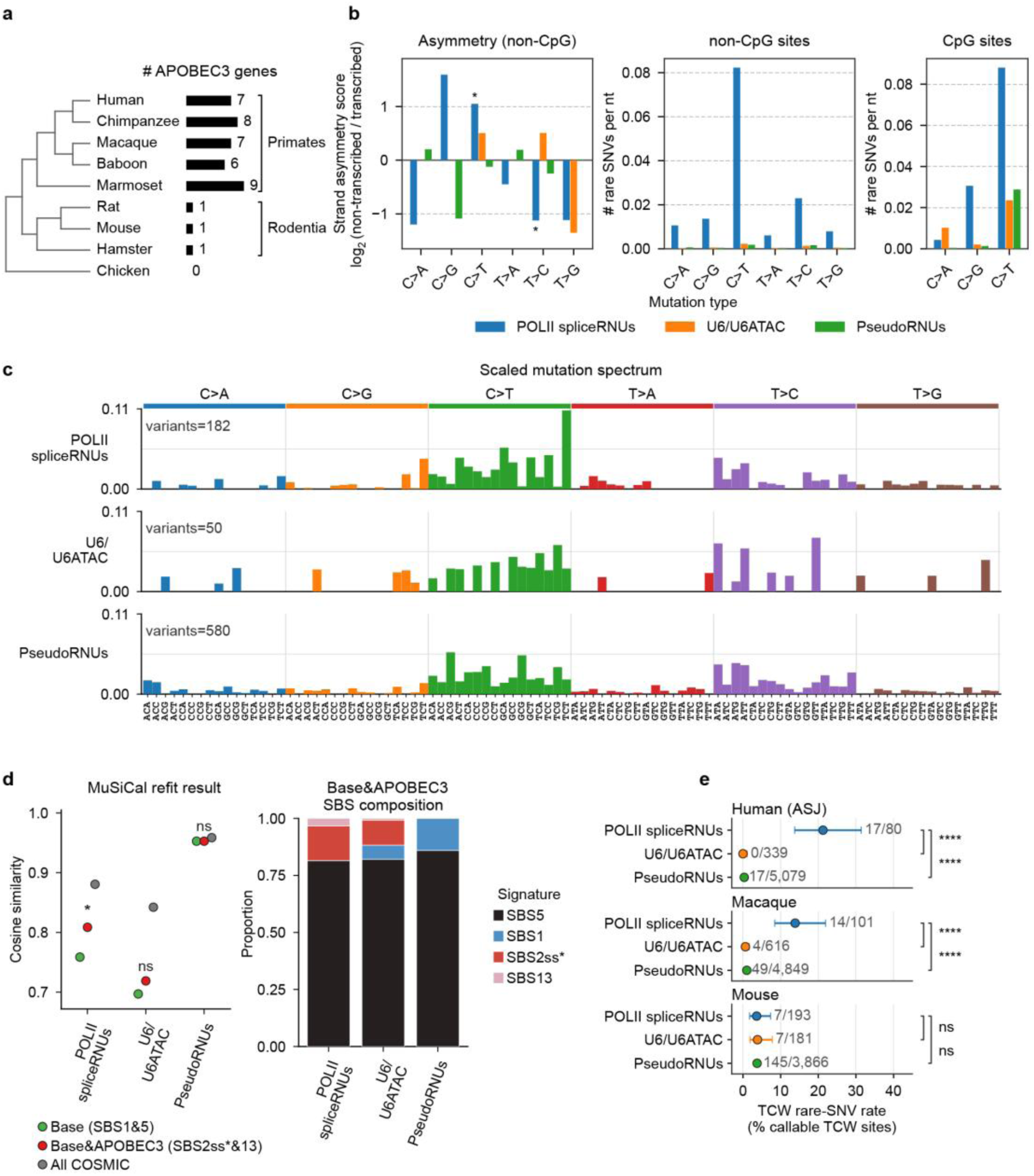
*Pol II spliceRNUs* exhibit mutational features compatible with APOBEC3 activity. **a**, APOBEC3 gene repertoire sizes are shown beside a simplified phylogenetic tree of representative species. **b**, Corrected rare SNV densities (1-mer) and transcriptional asymmetry scores for each gene group in the gnomAD v4 Ashkenazi Jewish (ASJ) population. CpG and non-CpG sites are shown separately, and asymmetry scores were calculated from non-CpG sites. Asterisks indicate *P* < 0.05 by two-sided Fisher’s exact tests. **c**, Scaled mutation spectra for *Pol II spliceRNUs*, U6/U6ATAC genes and *RNU* pseudogenes after normalization to genome-wide 3-mer compositions. The y axis shows the proportion of mutations in each 3-mer context. **d**, MuSiCal refitting under three signature sets (left) and fitted SBS contributions under the Base&APOBEC3 model (right). Asterisks indicate unadjusted empirical *P* < 0.05 from 1,000 targeted permutations; ns, not significant. **e**, Rare SNV rates at TCW motif sites in *Pol II spliceRNUs*, U6/U6ATAC genes and *RNU* pseudogenes in human ASJ, macaque and mouse. Points show observed rates; error bars, 95% Wilson confidence intervals. Brackets show *P* values from two-sided Fisher’s exact tests for *Pol II spliceRNUs* relative to each control set. ns: not significant; *: 0.01 < *P* ≤ 0.05; **: 0.001 < *P* ≤ 0.01; ***: 0.0001 < *P* ≤ 0.001; ****: *P* ≤ 0.0001.

We first examined transcriptional strand asymmetry. We analyzed CpG and non-CpG sites separately to limit confounding by C>T substitutions arising from spontaneous deamination of 5-methylcytosine at CpG sites^37^. At non-CpG sites, the C>T/G>A rare SNV density in *Pol II spliceRNUs* was 37.03- and 46.19-fold higher than in U6/U6ATAC genes and *RNU* pseudogenes, respectively (**Fig. 2b**; **Supplementary Table 10**). C>T substitutions were biased toward the non-transcribed strand (two-sided Fisher’s exact test, *P* = 0.002), in the same direction for C>G substitutions with marginal significance (*P* = 0.062; **Fig. 2b**), and similar strand bias was observed across four additional human datasets (**Supplementary Fig. 8-11**; **Supplementary Table 10 and 11**). This pattern is compatible with APOBEC3 activity on transcription-associated ssDNA and resembles the cytidine-biased mutagenesis reported at highly transcribed tRNA genes^38^.

We next asked whether the broader mutation spectrum carried APOBEC3-associated signatures. After normalizing rare SNV spectra to the genome-wide 3-mer composition (**Fig. 2c**), we used MuSiCal^39^ to fit combinations of mutational signatures. SBS1 and SBS5, prevalent clock-like components in normal human cells^40^, formed the baseline model. We then added SBS13 and SBS2ss*, an experimentally derived APOBEC3A-associated ssDNA damage spectrum distinct from COSMIC SBS2^35,36,41^, and compared this APOBEC3-augmented model with the full COSMIC reference set^42^. In *Pol II spliceRNUs*, cosine similarity increased from 0.759 to 0.809 after adding the APOBEC3-associated signatures. This improvement exceeded that obtained by adding two randomly selected alternative COSMIC signatures in 1,000 permutations (*P* = 0.042), and SBS2ss* plus SBS13 accounted for 18.5% of fitted activity (**Fig. 2d**; **Supplementary Table 12a**). No significant improvement was observed for U6/U6ATAC genes or *RNU* pseudogenes. This pattern was replicated across four additional human datasets, with APOBEC3-associated signatures accounting for 17.2–19.5% of fitted activity (**Supplementary Fig. 8-11**; **Supplementary Table 12b-d**). Because some APOBEC3-induced lesions may be assigned to SBS5 after repair^43,44^, these fitted proportions likely underestimate APOBEC3-attributable mutagenesis.

Because APOBEC3A and APOBEC3B preferentially deaminate cytosines in TCW motifs (where W denotes A or T), we next compared rare SNV densities at these TCW sites across gene groups within human, macaque and mouse (**Fig. 2e**; **Supplementary Table 13**). In ASJ *Pol II spliceRNUs*, 17 of 80 callable TCW sites carried rare variants (21.25%), compared with none in U6/U6ATAC genes (0/339) and 17 of 5079 sites in *RNU* pseudogenes (0.33%; 63.49-fold difference from *Pol II spliceRNUs*). Macaque showed a similar enrichment, with a TCW variant density of 13.86% (14/101) in *Pol II spliceRNUs*, 21.35- and 13.72-fold higher than in U6/U6ATAC genes and *RNU* pseudogenes, respectively (**Supplementary Table 13**). In contrast, TCW variant densities were similar across all three groups in mouse (3.63–3.87%; *P* = 1.00), indicating primate-specific enrichment of mutations at APOBEC3-preferred motifs.

Together, transcriptional strand asymmetry, reproducible improvement in APOBEC3-associated signature fits, and primate-biased TCW enrichment support a substantial APOBEC3-compatible component of *Pol II spliceRNU* hypermutability.

### Functional genomic evidence supports APOBEC3 engagement at hypermutable *Pol II spliceRNUs*

To obtain complementary molecular evidence linking APOBEC3 activity to *Pol II spliceRNUs*, we analyzed publicly available functional genomic datasets (**Supplementary Table 14**)^20,45,46^. We focused on APOBEC3B (A3B)-associated ssDNAs, the substrate for cytosine deamination^16^ and on R-loops, which expose non-template ssDNA and have been linked to elevated mutagenesis at functional snRNA loci^4^.

Using Flag-A3B SPI-seq data in human T47D cells^20^, we found that A3B-associated ssDNA was enriched 2.42- and 2.63-fold at *Pol II spliceRNUs* relative to U6/U6ATAC genes and *RNU* pseudogenes, respectively (**Fig. 3a, b**). Metagene profiles further showed that A3B-associated ssDNA increased near the transcription start sites, remained elevated across *Pol II spliceRNU* gene bodies and declined into flanking regions (**Fig. 3b**). These data support preferential A3B engagement with exposed ssDNA at human *Pol II spliceRNUs*.

**Fig. 3:**
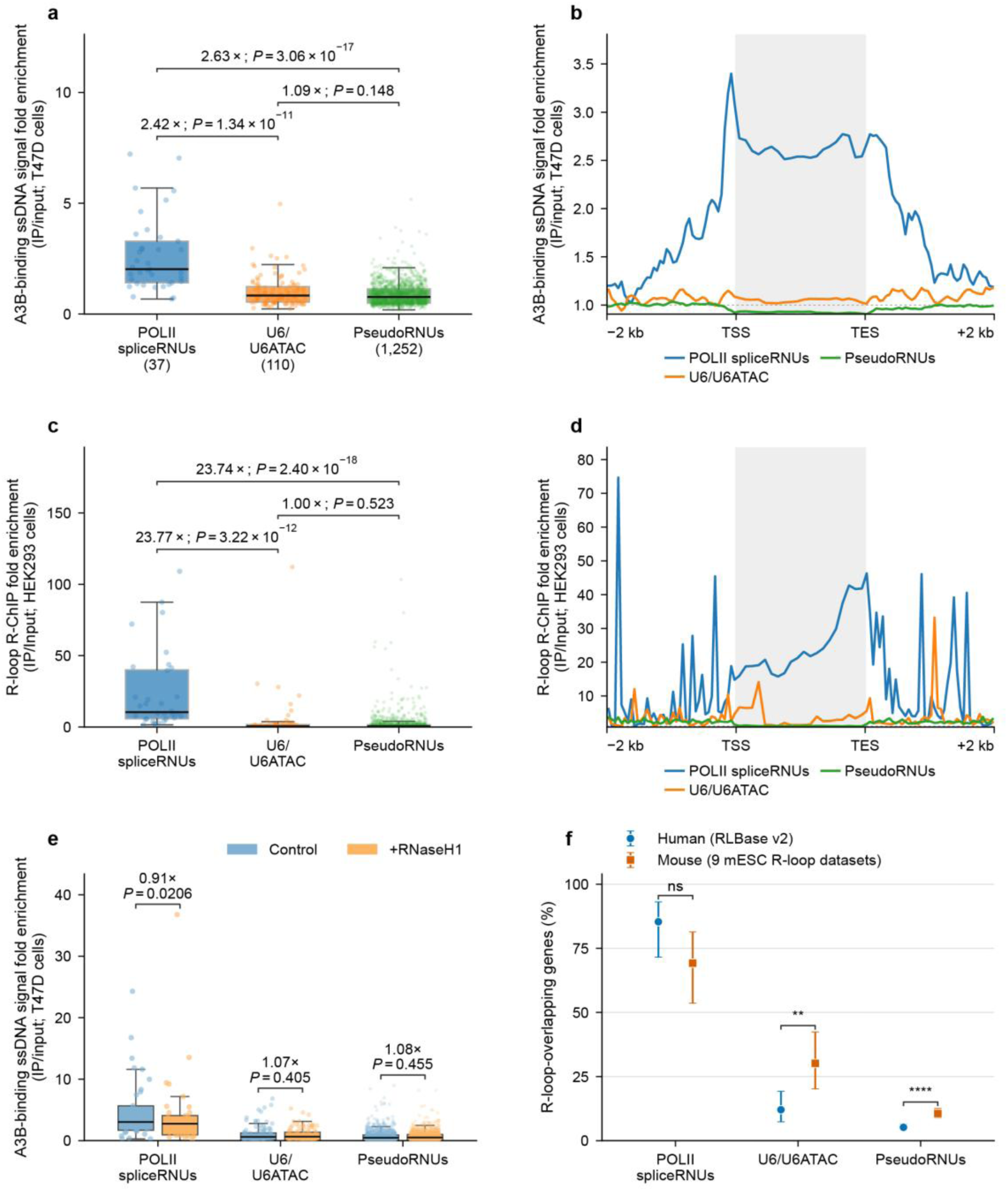
*Pol II spliceRNU* genes show A3B-associated binding and R-loop signals. **a,** Gene-body Flag-A3B SPI-seq fold enrichment in T47D cells. **b,** A3B-associated ssDNA SPI-seq profiles across 2-kb flanks and 1-kb scaled gene bodies. Curves show gene-level mean IP/Input signal from two siNT/DMSO datasets. TSS, transcription start site; TES, transcription end site. **c,** Gene-body RNase H1 D210N R-ChIP IP/Input fold enrichment in HEK293 cells. **d**, R-ChIP fold-enrichment profiles for the same gene groups. **e,** Gene-body Flag-A3B SPI-seq fold enrichment in control and RNase H1-overexpressing T47D cells. **f**, Fractions of genes overlapping human RLBase regions or the union of nine public mouse embryonic stem-cell R-loop peak sets. In **a**, **c** and **e**, boxes show medians and interquartile ranges, whiskers extend to 1.5 × IQR and outliers are not shown. Bracketed values are ratios of group medians. Comparisons in **a** and **c** used two-sided Mann-Whitney *U* tests; comparisons in **e** used paired two-sided Wilcoxon signed-rank tests. In **f,** points show proportions and error bars, 95% Wilson confidence intervals; species comparisons used two-sided Fisher’s exact tests. ns: not significant; *: 0.01 < *P* ≤ 0.05; **: 0.001 < *P* ≤ 0.01; ***: 0.0001 < *P* ≤ 0.001; ****: *P* ≤ 0.0001.

We next examined R-loop enrichment using published R-ChIP data from HEK293 cells^45^. *Pol II spliceRNUs* showed 23.77 and 23.74-fold higher gene-body R-loop enrichment than the comparison gene sets (**Fig. 3c, d**). To test the R-loop dependence of A3B engagement, we compared control and RNase H1-overexpressing T47D cells^20^. Although RNase H1 overexpression, reported to reduce R-loop signals by approximately 50%^47^, reduced the *Pol II spliceRNU* A3B-associated signal by only ∼9%, the residual signal remained 4.21–5.47-fold enriched over the comparison gene sets (**Fig. 3e**). Thus, R-loop reduction attenuates but does not account for the strong A3B-associated enrichment at *Pol II spliceRNUs*.

Comparing R-loop peak regions between human^46^ and mouse^48^ showed frequent overlap with *Pol II spliceRNUs* in both species (85.4% and 69.2%, respectively; **Fig. 3f**; **Supplementary Table 15**), with no significant species difference (two-sided Fisher’s exact test, *P* = 0.110). Thus, the much weaker hypermutability in mouse indicates that R-loop formation alone cannot explain the primate-specific *Pol II spliceRNU* hypermutability.

Together, these findings provide molecular support for APOBEC3 involvement in *Pol II spliceRNU* hypermutability. R-loops likely contribute by exposing ssDNA, but the lineage bias of the mutational patterns is better explained by APOBEC3 engagement than by R-loop abundance alone.

## Discussion

This study identifies Pol II-transcribed *spliceRNUs* as the principal hypermutable spliceosomal snRNA class and supports APOBEC3 activity as a major contributor. Transcriptional strand asymmetry and preferred motifs of mutations, APOBEC3-associated signatures and strong A3B-associated ssDNA enrichment all point to mutagenesis acting on exposed ssDNA^16,18,36^. This interpretation extends the recent observation that R-loop-forming regions, including snRNA genes, carry excess DNMs^4^: However, RNase H1 overexpression only partially reduces the A3B-associated signal, and mouse *Pol II spliceRNUs* frequently form R-loops despite much weaker hypermutability, indicating that R-loops alone are insufficient to explain the phenotype. Together, these findings suggest that R-loops provide susceptible ssDNA substrates, while APOBEC3 engagement and subsequent lesion processing determine mutational outcome^17,20,36^.

The cross-species patterns further strengthen this interpretation. The stronger relative excess in primates than in rodents and chicken is consistent with lineage-dependent APOBEC3 activity. APOBEC3 genes have expanded and diversified in primates, likely driven in part by host–virus and host–retroelements conflicts^21,30^, whereas mouse and other rodents carry a single *Apobec3* and can restrict retroviruses through mechanisms that do not require strong DNA deamination^49,50^. Such immune-system innovations may therefore have had collateral effects on host-genome mutagenesis. Consistent with this possibility, our recent work identified AID/APOBEC off-target activity as a major contributor to the germline hypermutability of replication-dependent histone genes^51^. Additionally, the weaker *Pol II spliceRNU* hypermutability observed in chicken, an avian outgroup lacking APOBEC3, extends this contrast beyond mammals and is consistent with APOBEC3 contributing to the stronger primate signal. Together, these findings support off-target mutagenesis at susceptible host loci with important functions as an evolutionary cost of cytidine-deaminase-based genomic defense. Broader comparisons across mammals and other vertebrates will help define the evolutionary extent of this trade-off.

A key limitation is that the germline mutation data and A3B/R-loop functional datasets come from different biological systems. The functional genomic data localize A3B-associated ssDNA and R-loop signals to these loci but do not identify the specific APOBEC3 enzymes, germline cell types or developmental stages responsible for heritable mutations. Direct perturbation of candidate APOBEC3 enzymes in germline-relevant systems, coupled with measurements of R-loops, ssDNA exposure and newly acquired mutations, will be required to establish enzyme-specific causality and quantify the relative contributions of R-loop-associated and other transcription-generated substrates.

## Methods

### Data preparation

#### Population variant data

For wild mice and rats, no suitable precomputed variant callset was available. We therefore compiled population-scale whole-genome sequencing data and implemented a standardized variant-calling and filtering pipeline (metadata provided in **Supplementary Table 16**).

Raw data were downloaded from the Sequence Read Archive using parallel-fastq-dump (https://github.com/rvalieris/parallel-fastq-dump) and converted to FASTQ format. Reads were processed with fastp (v0.24.0)^52^ to remove adapter sequences and trim low-quality bases, producing clean reads for downstream analyses. Clean reads were aligned to the reference genome using BWA-MEM2 (v2.2.1)^53^. PCR duplicates were marked using MarkDuplicates in GATK (v4.6.1.0)^54^, and variants were called per sample using GATK HaplotypeCaller in GVCF mode to generate per-sample gVCF files. Per-base depths across the whole reference genome and individual-level average sequencing depth metrics were subsequently computed using mosdepth (v0.3.10)^55^ with default parameters. Read alignment, preprocessing and variant calling were implemented using nf-core/sarek (v3.7.0)^56^. Per-sample gVCFs were subsequently jointly genotyped and merged using GATK to produce a multi-sample VCF for each species.

Cryptic relatedness can confound the analysis with rare variants. After filtering and pruning variants based on linkage disequilibrium, we estimated within-species pairwise relatedness score using Hail’s pc_relate module^57^. Pairs with relatedness score > 0.05 were considered related. We then used Hail’s maximal_independent_set module to remove the minimum number of samples required to obtain a dataset with unrelated individuals. For variant sites, we performed hard filtering using GATK4 VariantFiltration on SNPs and INDELs independently, excluding SNPs with “QUAL<30, QD<2, FS>60, MQ<40, MQRankSum<-12.5, ReadPosRankSum<-8 or SOR>3”^58^. To ensure accurate estimation of AF, genotypes with genotype quality (GQ) < 20 were masked as ‘./.’ and excluded from AF calculations.

For human, chimpanzee, macaque, baboon and chicken, we directly used publicly available variant datasets (**Supplementary Table 16**). These datasets were processed using the same individual-level and site-level filtering strategy described above to ensure comparability across species.

The primary human analysis used 1,736 individuals from the gnomAD v4 Ashkenazi Jewish (ASJ) group. The quality-filtered variant datasets used for the cross-species comparisons were derived from populations of 198 chimpanzees, 514 macaques, 110 baboons, 148 mice, 89 rats and 191 chickens. AF was calculated as the corresponding allele count divided by the number of called alleles. Rare variants were defined using inclusive AF thresholds of ≤ 1 × 10^-3^ for human, ≤ 1 × 10^-2^ for chimpanzee, ≤ 2 × 10^-3^ for macaque, ≤ 2 × 10^-2^ for baboon, ≤ 1 × 10^-2^ for mouse, ≤ 2 × 10^-2^ for rat and ≤ 1 × 10^-2^ for chicken. Species-specific AF thresholds were chosen to accommodate differences in cohort size.

#### Replication across human populations and downsampling analysis

We repeated the human analysis in the gnomAD v4 East Asian (EAS; 2,598 individuals), South Asian (SAS; 2,417 individuals) and Finnish (FIN; 5,316 individuals) groups, and in a fixed 2,000-individual randomly selected subset of the gnomAD v3.1.2 Human Genome Diversity Project and 1000 Genomes Project dataset (HGDP+1KG; random2000). The random2000 set was all drawn from the unrelated individuals.

To assess sensitivity to sample size and the species-specific rarity thresholds, we independently sampled 500, 200 and 100 individuals without replacement from the pool of 2,000 unrelated HGDP+1KG individuals, drawing one subset per sample size. Allele frequencies were recalculated from the selected genotypes, with AF thresholds of ≤ 2 × 10^-3^, ≤ 1 × 10^-2^ and ≤ 2 × 10^-2^, respectively, matching the macaque, mouse and rat analyses. Rare SNV counts, masked positions and effective callable lengths were recalculated for the fixed *RNU* and control intervals.

#### Human de novo mutation data

DNMs from 9,652 trios were obtained from Pálsson et al.^28^. We restricted analyses to 618,192 *de novo* SNVs and used these variants to estimate the genome-wide mutation spectrum and de novo mutation rates across different gene groups. For each gene group, the total number of callable nucleotides was calculated as the sum of filtered region lengths across genes, multiplied by the number of trios and by two to account for diploidy. The mutation rate (per site per generation) was then estimated as the number of DNMs divided by the total callable nucleotide count.

#### Excluded genomic regions

To minimize potential biases in downstream analyses, we excluded genomic regions with low mappability or abnormal sequencing coverage. Genomic mappability for each species was calculated using GenMap (v1.3.0)^59^ with parameters ‘-K 150 -E 1’. Per-base coverage of the human genome was obtained from gnomAD; for other species, coverage was estimated from a set of randomly selected sequenced individuals using deepTools bamCoverage (v3.5.6)^60^. Regions with mappability < 1 or coverage outside 0.5-1.5× the mean sequencing depth were excluded from variant density calculation and other related analyses.

### Identification and classification of snRNA genes

Within each species, loci were assigned to three mutually exclusive classes: *Pol II spliceRNU*, U6/U6ATAC genes and *RNU* pseudogenes. Intergenic regions were included as genomic controls. The functional genomics datasets used for snRNA gene identification and classification are listed in **Supplementary Table 1**.

#### Human

Human gene coordinates were obtained from GENCODE v44^61^. We defined the human *Pol II spliceRNU* set by intersecting POLR2A-supported loci with a previously reported catalogue of 200 putatively functional snRNA genes^6^. Following the chromatin-based approach of Kosmyna et al.^13^, we analyzed ENCODE POLR2A ChIP-seq data from A549, H1, HeLa-S3, HepG2, K562 and MCF-7 cells. A locus was considered POLR2A-supported if it showed mean POLR2A enrichment >2-fold and overlapped the corresponding consensus POLR2A peak in at least one cell line. Of 49 POLR2A-supported loci, 44 overlapped the curated 200-locus snRNA catalogue. Three U1-family loci (*RNVU1-26*, *RNVU1-2A* and *RNVU1-28*) contained no uniquely mappable callable sequence and were excluded, leaving 41 *Pol II spliceRNUs* for downstream analyses (**Supplementary Fig. 1a)**. Of the 118 previously reported putatively functional U6/U6ATAC genes^6^, 116 contained callable sequence and were retained. The remaining eligible autosomal *RNUs* comprised an initial pseudogene set of 1,559 loci, of which 1,362 passed the length and effective callable base filters (**Supplementary Fig. 1**; **Supplementary Table 2**).

#### Macaque

Macaque snRNA genes were obtained from Ensembl Mmul_10 release 115 on the rheMac10 assembly. Initial functional annotation used cCREs derived from two published single-cell chromatin-accessibility studies of the macaque brain^62,63^. We intersected the cCRE intervals from the two studies to define a consensus set of candidate regulatory regions. For each annotated snRNA gene, we defined a promoter-proximal interval extending 50 bp upstream and downstream of its strand-specific transcription start site (TSS). Genes whose TSS ±50-bp interval overlapped a consensus cCRE were classified as putatively functional, whereas genes without such overlap were assigned to an operational *RNU* pseudogene set. Among the cCRE-supported loci, genes annotated as U6 or U6atac constituted the U6/U6ATAC set.

The *Pol II spliceRNU* set was defined by syntenic correspondence to the 41 human *Pol II spliceRNUs*. Human *Pol II spliceRNU* gene intervals were projected onto rheMac10 using the UCSC hg38-to-rheMac10 chain and liftOver, with a minimum mapped fraction of 0.20. Gene-body assignments required a same-family macaque snRNA annotation to overlap at least 30% of both the projected human interval and the macaque locus. For genes lacking a gene-body assignment, flanking intervals were mapped independently with a minimum mapped fraction of 0.50, beginning with anchors 200–1,000 bp outside the human locus. Same-family macaque candidates within 100 kb of the projected position were assessed by proximity and BLASTN-short sequence similarity. One-to-one syntenic assignments were retained only when a macaque locus mapped uniquely to a single human locus. U6/U6ATAC and pseudogene membership otherwise followed the cCRE-based classification. After the callable-sequence filters described above, the macaque analysis included 28 *Pol II spliceRNUs*, 193 U6/U6ATAC genes and 1,215 *RNU* pseudogenes (**Supplementary Fig. 1c, d**; **Supplementary Table 4**).

#### Chimpanzee

We applied the synteny-based procedure used for macaque *Pol II spliceRNUs* separately to the three human-reference *RNU* classes in chimpanzee: 41 *Pol II spliceRNUs*, 116 U6/U6ATAC genes and 1,362 *RNU* pseudogenes (**Supplementary Fig. 4a**). Target loci were Ensembl release 115 snRNA annotations on panTro5. Core hg38 intervals were projected by liftOver with a minimum mapped fraction of 0.20. Family-compatible candidates required overlap covering at least 30% of both projected and annotated intervals; candidates within 0.05 of the best overlap fractions were retained for ambiguity assessment. Where core assignment failed, independent flanking intervals and subsequently extended locus windows were mapped with a minimum mapped fraction of 0.50. Same-family annotations within 100 kb of the predicted position were evaluated by proximity, prioritizing candidates with BLASTN-short identity ≥65% and query coverage ≥60%; the nearest annotation could be selected when sequence support was absent. Single-target assignments were retained only when the target was not reused within the same input class. After assignment, four pseudogenes sharing targets with retained *Pol II spliceRNUs* or U6/U6ATACs were excluded to maintain distinct target sets. This yielded 28 *Pol II spliceRNUs*, 110 U6/U6ATACs and 1,149 *RNU* pseudogenes (**Supplementary Fig. 4b**; **Supplementary Table 3**). Variant analysis used their original GRCh38 coordinates and human-reference class labels.

#### Baboon

We extracted Ensembl release 115 Panubis1.0 snRNA genes and retained 1,681 autosomal annotations. Human *Pol II spliceRNUs* and U6/U6ATAC sets were transferred using the tiered positional assignment applied to macaque. The forward chain was generated by swapping the UCSC Panubis1.0-to-GRCh38 chain. Core projections used a minimum mapped fraction of 0.20 and required family-compatible annotations with ≥30% overlap of both intervals; candidates within 0.05 of the best overlap fractions were retained. Where core assignment failed, flanking anchors or extended intervals were mapped at a minimum fraction of 0.50. Same-family annotations within 100 kb of the predicted centre were considered, prioritizing nearby candidates supported by ≥65% sequence identity and ≥60% query coverage; the nearest annotation could be selected when sequence support was absent. Unique assignments without target reuse yielded 18 *Pol II spliceRNUs* and 84 U6/U6ATAC genes, after callable-sequence filters. The remaining 1,307 autosomal snRNA annotations formed the pseudogene set (**Supplementary Fig. 4c, d**; **Supplementary Table 5**).

#### Mouse

Mouse RNUs were reannotated using regulatory annotation and phylogenetic relationships (**Supplementary Fig. 5a**). snRNA loci were obtained from GENCODE vM25 on the mm10 assembly. Candidate functional loci were identified by overlap with mouse cCRE annotations, with non-spliceosomal RNUs excluded. Mouse candidate sequences were aligned with annotated human snRNAs using MAFFT^64^ with local-pair alignment and 1,000 refinement iterations and trimmed using trimAl^65^ with the automated option. Maximum-likelihood phylogenies were inferred using IQ-TREE (v2.3.5)^66^ and 1,000 ultrafast bootstrap replicates; Bayesian information criterion-based model selection identified HKY+F+R2. Phylogenetic assignment relative to human snRNAs was used to distinguish *Pol II spliceRNUs* from U6/U6ATAC genes. The initial annotation comprised 50 *Pol II spliceRNU* candidates, 64 U6/U6ATAC genes and 1,264 *RNU* pseudogenes; callable-sequence filtering retained 39, 63 and 1,032 loci, respectively. Of the 11 excluded Pol II candidates, seven lacked callable sequence and four contained fewer than 50 effective callable bases. All 39 retained *Pol II spliceRNUs* overlapped the available mm10 cCRE annotation (**Supplementary Fig. 5b**; **Supplementary Table 6**).

#### Rat

Rat loci were drawn from Ensembl release 112 mRatBN7.2 (rn7) snRNA annotations. Family assignments used Rfam^67^ covariance models with Infernal (v1.1.5)^68^; clear Ensembl/RGD family annotations were retained when no model passed. U7 loci and unresolved or conflicting family assignments were excluded. Promoter accessibility was assessed using published multi-organ and nervous-system ATAC-seq atlases^69,70^. Peaks were transferred from rn6 to rn7 with a minimum mapped fraction of 0.95. For the *Pol II spliceRNU* and U6/U6ATAC sets, the annotated TSS ±50 bp had to overlap the genomic intersection of peak intervals from the two atlases. This identified 46 *Pol II spliceRNUs* and 37 U6/U6ATAC genes after callable filtering (**Supplementary Fig. 5c, d**; **Supplementary Table 7**). The *RNU* pseudogene set comprised 847 autosomal loci lacking ATAC-seq support in either atlas. We excluded loci whose promoter intervals could not be mapped to rn6 and those with reciprocal positional correspondence to mouse functional *Pol II spliceRNUs* or U6/U6ATAC genes despite lacking rat ATAC-seq support. Mouse–rat correspondence used bidirectional liftOver with a minimum mapped fraction of 0.50, concordant strand and ≥30% overlap of both intervals.

#### Chicken

Chicken snRNA loci were obtained from Ensembl release 115 on GRCg7b and restricted to autosomal non-U7 annotations. Regulatory support was evaluated using chromatin accessibility, active chromatin states and active histone marks^71^. Accessibility and chromatin-state annotations were assessed within TSS ±50 bp; H3K27ac and H3K4me3 were assessed within TSS ±500 bp. Peak intervals shared between biological replicates were retained, and GRCg6a annotations were transferred to GRCg7b with a minimum mapped fraction of 0.95. *Pol II spliceRNUs* required a compatible snRNA-family annotation and either strand-specific nascent transcription supported by at least one chromatin evidence category, or Pol II Ser5P occupancy accompanied by accessible chromatin^72^. U6/U6ATAC loci required annotation and chromatin support. Remaining autosomal non-U7 snRNAs formed the operational pseudogene comparator set. The initial annotation comprised 15 *Pol II spliceRNU* candidates, 9 U6/U6ATAC genes and 48 *RNU* pseudogenes; callable-sequence filtering retained 14, 9 and 38 loci, respectively (**Supplementary Fig. 6b, c**; **Supplementary Table 8**).

#### Intergenic control loci

For each species, we sampled 1,000 non-overlapping 150-bp autosomal intergenic control intervals, allocated among chromosomes in proportion to chromosome length following D’Souza et al.^12^. Intervals overlapping annotated gene bodies or their 10-kb flanking sequences, assembly gaps or other sampled intervals were excluded. From a pool of 10,000 candidates, we selected without replacement 1,000 intervals for which all 150 bases were callable. All species controls were restricted to assembled autosomes. The same human control coordinates were used across ancestry groups and random subsets. Intergenic controls were subjected to the same variant-position masking as *RNU* loci.

### Rare SNV density and enrichment relative to controls

For each gene or control locus g in dataset s, we calculated

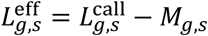

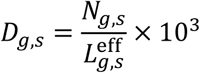

where *L*^call^ is the callable length before variant masking, *M* is the number of uniquely masked callable positions and *N* is the retained rare alternative allele count. Density was expressed as rare SNVs per effective callable kilobase. Loci with a positive eligible denominator but no qualifying alternative allele entries were assigned zero density and retained in all summaries and comparisons.

For each class *k*, enrichment relative to controls (**Fig. 1b**) was calculated from unweighted arithmetic means:

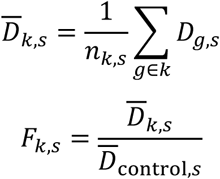

The control ratio was therefore one by definition within each species or human cohort. Pairwise mean-density fold differences were calculated analogously. The same gene identities were used when comparing human ancestry groups and random subsets.

### Transcriptional strand asymmetry and variant density

Mutation densities were corrected at the trinucleotide-channel level before aggregation (**Fig. 2b**). For mutation type channel *i*, let *N_i_* denote the reference-context count, *m_i_* the number of masked records and *n_i_* the retained rare SNV count. We calculated the effective number of sites *D_i_*, Poisson rate ℎ*_i_* and corrected mutation amount *u_i_* as

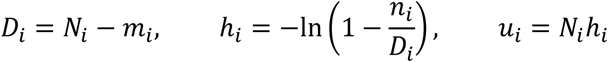

For each central-base substitution, corrected amounts were summed across flanking contexts and divided by the corresponding total number of reference-context counts. Trinucleotides containing a CpG dinucleotide were analyzed separately. Reverse-complementary substitutions were pooled for the density plots, such that, for example, C>T includes G>A.

Strand asymmetry was evaluated at non-CpG sites. For a pyrimidine substitution *s* and its reverse-complementary partner *̄S* after orienting sequences by transcriptional strand, the score was 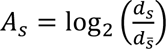, *d_S_* is the corrected density of the displayed C- or T-based substitution and *d_̄S_* is the density of its G- or A-based partner. For example, the C>T score is log_2_(*d_C_*_>*T*_/*d_G_*_>*A*_) . Positive scores indicate enrichment of the pyrimidine substitution on the non-transcribed relative to the transcribed strand. Two-sided Fisher’s exact tests compared the rows (*n*_NTS_, *D*_NTS_ − *n*_NTS_) and (*n*_TS_, *D*_TS_ − *n*_TS_), where *D_s_* = ∑*_i_*_∈_*_s_* (*N_i_* − *m_i_*) . Thus, the asymmetry score used corrected densities, whereas its significance test used observed counts.

For the accompanying density panels, reverse-complementary substitutions were combined using the same reference-composition-weighted Poisson-rate correction. CpG and non-CpG channels were analyzed separately, with CpG channels defined by the presence of a CG dinucleotide in the reference trinucleotide. Population-replication analyses in **Supplementary Fig. 8-11** used the same corrected-density asymmetry score. As in the ASJ analysis, Fisher’s exact tests in these panels were based on observed, uncorrected variant counts.

### Mutation signature analysis

We refitted the observed 96-channel substitution spectra using reference single-base substitution (SBS) signatures^42^ to deconvolute the contributions of various mutational processes in *Pol II spliceRNUs*. Prior to signature decomposition, we computed the 3-mer frequencies for all usable genomic regions to establish a genome-wide background. The mutational catalog of each gene group was normalized to this background composition to account for base composition biases. Because APOBEC3 enzymes preferentially deaminate cytosines in ssDNA, we replaced standard SBS2 with the experimentally derived ssDNA-associated SBS2ss* profile from Liu et al.^41^

To correct for potential saturation effects from highly mutable sites, we applied the same Poisson-based adjustment detailed in the ’Transcriptional strand asymmetry and variant density’ section to the mutational counts before signature refitting. Deconvolution was performed using MuSiCal (v1.0.0)^39^, which employs a likelihood-based sparse non-negative least squares algorithm. We set a likelihood threshold of 0.005 to ensure robust signature assignment and limit false-positive identifications. Signatures were grouped etiologically based on COSMIC annotations: SBS2ss* and SBS13 were classified as APOBEC3-associated.

We assessed whether adding SBS2ss* and SBS13 improved reconstruction more than adding two alternative signatures using a permutation test. SBS1 and SBS5 were retained in every model. In each of 1,000 iterations, two distinct signatures were sampled from the remaining 63 signatures and the resulting alternative-signature model was refitted using the same settings. Sampling was without replacement within each pair; pairs could recur between iterations. The alternatives were sorted before sampling with Python’s pseudorandom generator, and the same sampled pair was evaluated for all three gene groups.

For each gene group, the one-sided empirical *P* value was the fraction of alternative fits with cosine similarity at least as high as the Base&APOBEC3 fit:

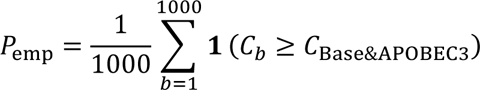

The same sampled pairs were used for the three gene groups. These analyses were performed separately in ASJ, EAS, SAS, FIN and random2000.

### TCW rare-variant rates across species

Population TCW analyses used TCA and TCT in transcript-oriented reference sequence, where W denotes A or T. For the all-substitution analysis in **Fig. 2e**, the numerator was the sum of C>A, C>G and C>T records at these contexts. The denominator was the TCA/TCT reference-context count minus the sum of masks for these substitutions. Variant burdens were expressed as percentages, with 95% Wilson confidence intervals. *Pol II spliceRNUs* were compared with U6/U6ATAC genes and *RNU* pseudogenes using two-sided Fisher’s exact tests.

### Gene sets for functional genomics analysis

Functional genomic analyses included the *Pol II spliceRNUs*, U6/U6ATAC genes and *RNU* pseudogenes defined above, together with expressed protein-coding genes identified from the fetal testis expression catalogue of Lecluze et al.^73^. Signals at *RNU* loci were quantified across annotated gene bodies. For protein-coding genes, coding sequence (CDS) intervals from all annotated transcripts of each gene were merged, with overlapping bases counted once. Gene identifiers were matched after removal of annotation-version suffixes. CDS coordinates were obtained from GENCODE v44 for GRCh38 and GENCODE v44lift37 for GRCh37.

For analyses using GRCh37-based signal tracks, GRCh38 *RNU* intervals were converted to GRCh37 coordinates using UCSC liftOver with the hg38-to-hg19 chain file and a minimum mapped fraction of 0.95. This yielded 37 *Pol II spliceRNUs*, 110 U6/U6ATAC genes and 1,252 *RNU* pseudogenes with successfully mapped intervals. The corresponding gene identifiers were used to define the matched *RNU* sets for GRCh38-based analyses.

### APOBEC3B-associated ssDNA enrichment

We obtained processed single-stranded DNA protein-immunoprecipitation sequencing (SPI-seq) tracks from T47D cells expressing inducible Flag–A3B^20^. Gene-level enrichment and metagene profiles were calculated from the two non-targeting siRNA/DMSO replicates in GSE193228 (GSM5777032 and GSM5777033; **Fig. 3a, b**). The GRCh37 bigWig tracks contained MACS2-derived fold-enrichment-over-input signals^74^.

At each callable position, positive enrichment values were averaged across available replicates. Gene-level enrichment was then calculated as the arithmetic mean of these position-level values across callable positions with available signal, such that each genomic position contributed equally regardless of whether one or both replicates provided a measurement. The same locus sets and position-level signal definition were used for gene-level comparisons and metagene profiles. Gene-level enrichment distributions between locus classes were compared using two-sided Mann–Whitney *U* tests with the asymptotic approximation and correction for ties. Fold differences between classes were calculated as ratios of median gene-level enrichment values.

To assess the effect of RNase H1 overexpression on A3B-associated SPI-seq enrichment, we analyzed an independent FLAG–A3B SPI-seq experiment from GSE193227^20^, comparing the lipid-transfection control (GSM5795929) with RNase H1-overexpressing cells (GSM5795927; **Fig. 3e**). For each condition, gene-level enrichment was calculated as the mean signal across callable positions with finite track values within *RNU* gene bodies or the merged CDS intervals of expressed protein-coding genes. Control and RNase H1 measurements were matched by gene identifier, and only genes with measurements in both conditions were retained. Differences between conditions were assessed within each gene class using two-sided Wilcoxon signed-rank tests. Retained A3B-associated ssDNA signal was expressed as the ratio of the median gene-level enrichment under RNase H1 overexpression to that in the control condition.

### R-loop enrichment measured by R-ChIP

R-loop signals were quantified using R-ChIP data from HEK293T cells expressing catalytically inactive RNase H1 (D210N) (GSE97072)^45^. For each locus, normalized coverage was averaged over callable positions in the *RNU* gene bodies or the merged CDS intervals of expressed protein-coding genes. R-ChIP enrichment was then calculated as the ratio of the mean normalized IP coverage across the three replicates to the corresponding mean normalized input coverage. The same ratio-of-means calculation was applied to each bin of the metagene profiles. Enrichment values were calculated only when the corresponding mean input coverage was greater than zero. The same signal definition was used for gene-level comparisons and metagene profiles. Gene-level enrichment distributions between locus classes were compared using two-sided Mann–Whitney *U* tests, and fold differences were calculated as ratios of median gene-level enrichment values (**Fig. 3c, d**).

### Cross-species comparison of R-loop overlap at *RNU* loci

Human R-loop overlap was assessed using the GRCh38 consensus R-loop regions distributed through RLBase^46^. For mouse, we constructed a composite R-loop annotation by taking the union of nine publicly available mouse embryonic stem cell peak sets^48^: DRIP-seq and DRIP-RNA-seq from GSE70189 and GSE67581; two S9.6 CUT&Tag datasets from GSE173103; two MapR and two BisMapR datasets from GSE160578; and the published HBD-seq consensus intervals from GSE254764. The first eight datasets were obtained as RLBase-processed mm10 peak files. Overlapping or directly adjacent intervals were merged to generate the final mouse R-loop union.

A gene was considered R-loop-overlapping if at least 1 bp of its annotated gene body intersected an R-loop interval. For human loci, the overlapping base was additionally required to fall within the callable mask, and only genes containing at least one callable base were included in the denominator. Cross-species comparisons used the human and mouse *RNU* gene sets defined in **Fig. 1b**. Overlap proportions and 95% Wilson confidence intervals were calculated separately for each species and gene class (**Fig. 3f**). Human and mouse overlap proportions were compared within each gene class using two-sided Fisher’s exact tests.

## Data availability

Summary data supporting the findings are included in the main text and supplementary materials. All the analyses were based on published data.

## Code availability

Code for the analyses performed in this study is available on GitHub at https://github.com/xuejinghub/RNU_mutation_rate_analysis.

## Acknowledgments

This work was supported by the National Natural Science Foundation of China (32470690).

## Supplementary Figure

**Supplementary Fig. 1.**
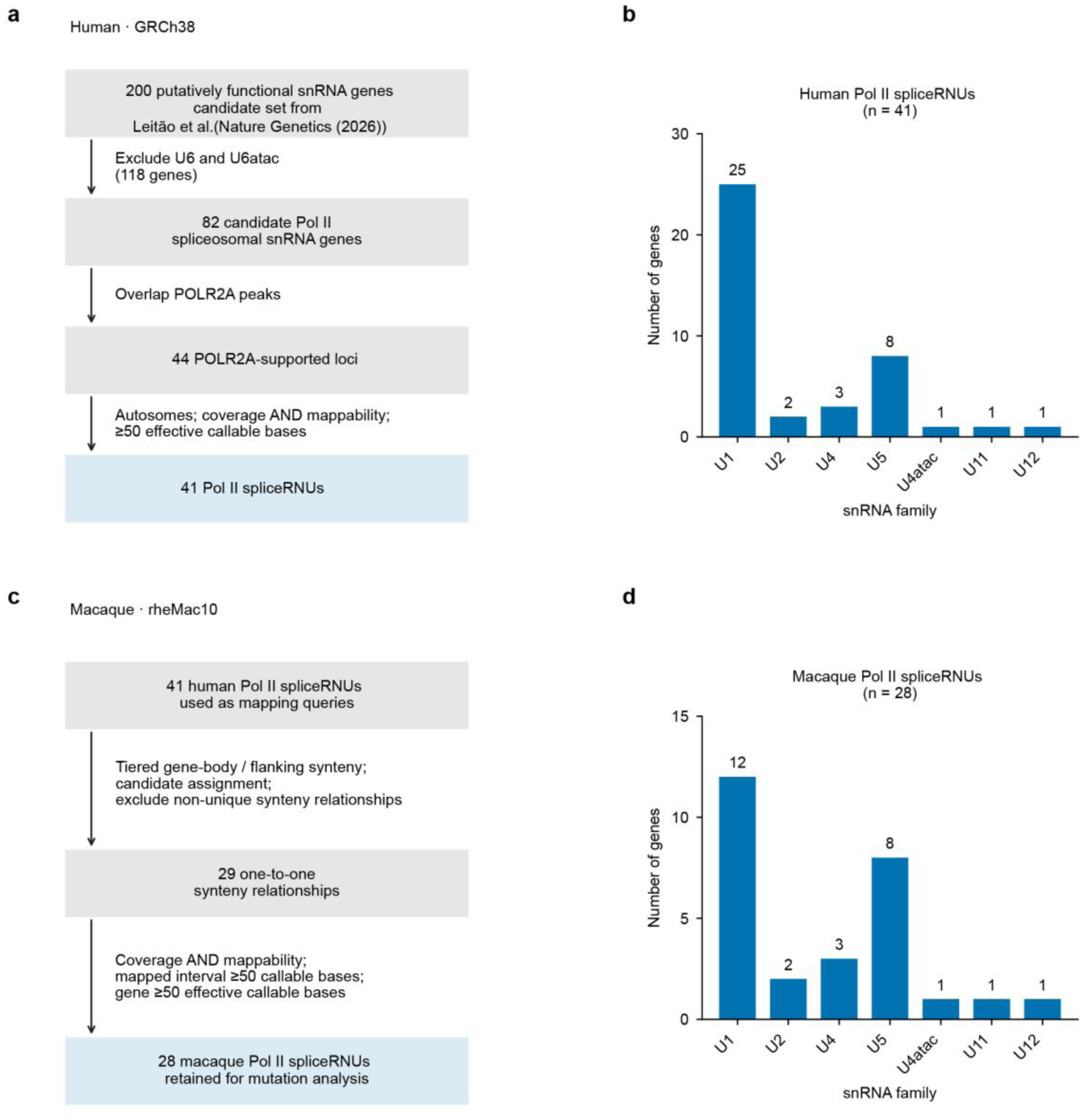
Definition and family composition of human and macaque *Pol II spliceRNU* analysis sets. **a**, Filtering strategy used to retain genes expressing possible human *Pol II spliceRNUs*. The workflow begins with 200 putatively functional spliceosomal snRNA candidates in the local catalogue, with the literature-based prioritization framework cited to Leitão et al. **b**, Number and distribution of annotated human *Pol II spliceRNUs*. **c**, Filtering strategy used to retain genes expressing possible macaque *Pol II spliceRNUs*. **d**, Number and distribution of annotated *Pol II spliceRNUs* of macaque.

**Supplementary Fig. 2.**
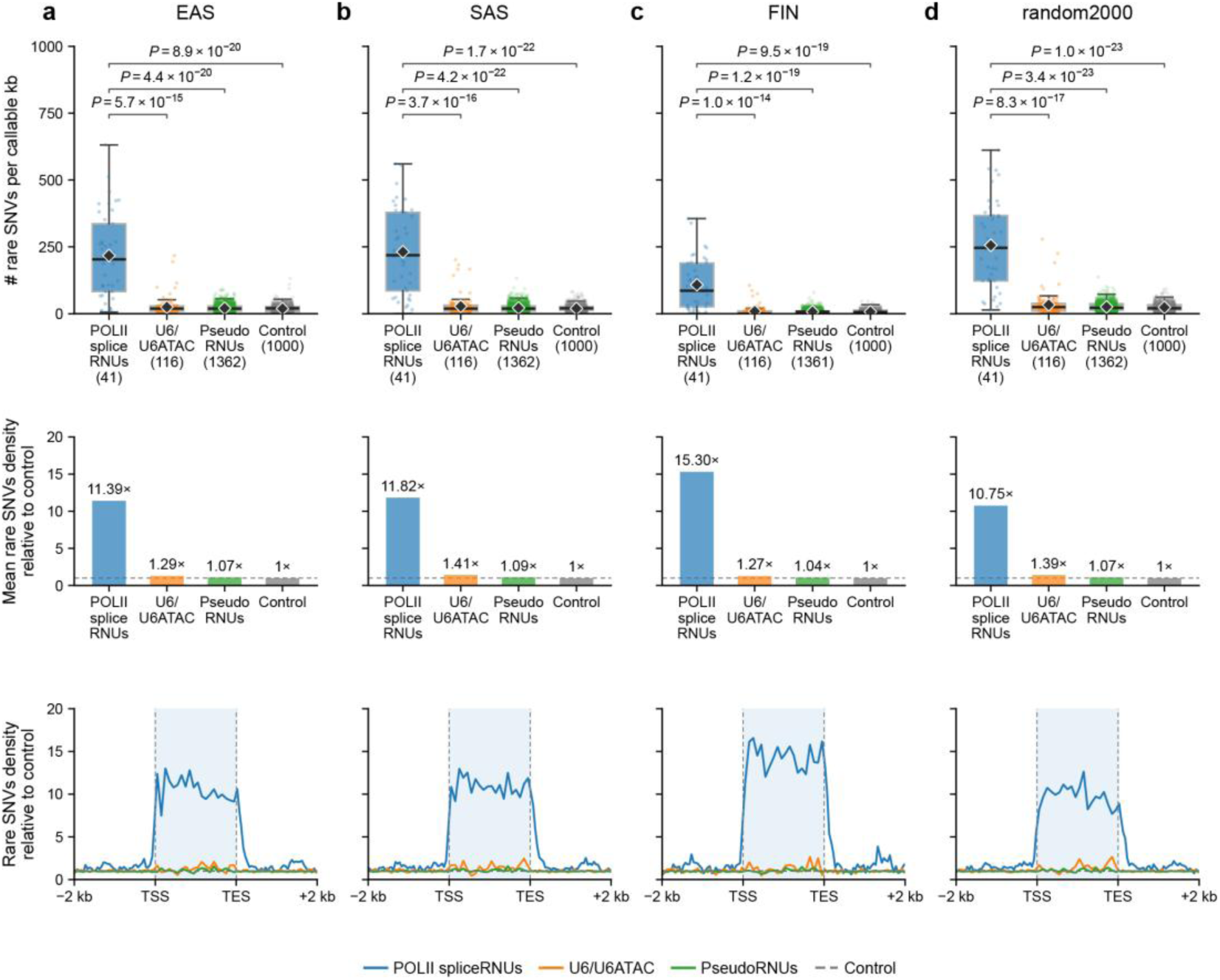
Rare-SNV enrichment and positional profiles across human cohorts. **a**–**d**, gnomAD v4 East Asian (EAS; **a**), South Asian (SAS; **b**) and Finnish (FIN; **c**) ancestry groups, and a fixed random 2,000-individual HGDP+1KG subset (random2000; **d**). All analyses use GRCh38 and AF ≤ 1 × 10^-3^. Top row, Gene- or locus-level rare SNV density for the same 41 *Pol II spliceRNUs*, 116 U6/U6ATAC genes, 1,362 *RNU* pseudogenes and 1,000 autosomal intergenic control regions in every cohort. Each human control is a 150-bp interval on autosome chromosomes. Boxes show the interquartile range, centre lines medians, whiskers the most extreme observations within 1.5 times the interquartile range, points individual loci and diamonds arithmetic means. Brackets show *P* values from two-sided asymptotic Mann–Whitney *U* tests, comparing *Pol II spliceRNUs* with each other class.

**Supplementary Fig. 3.**
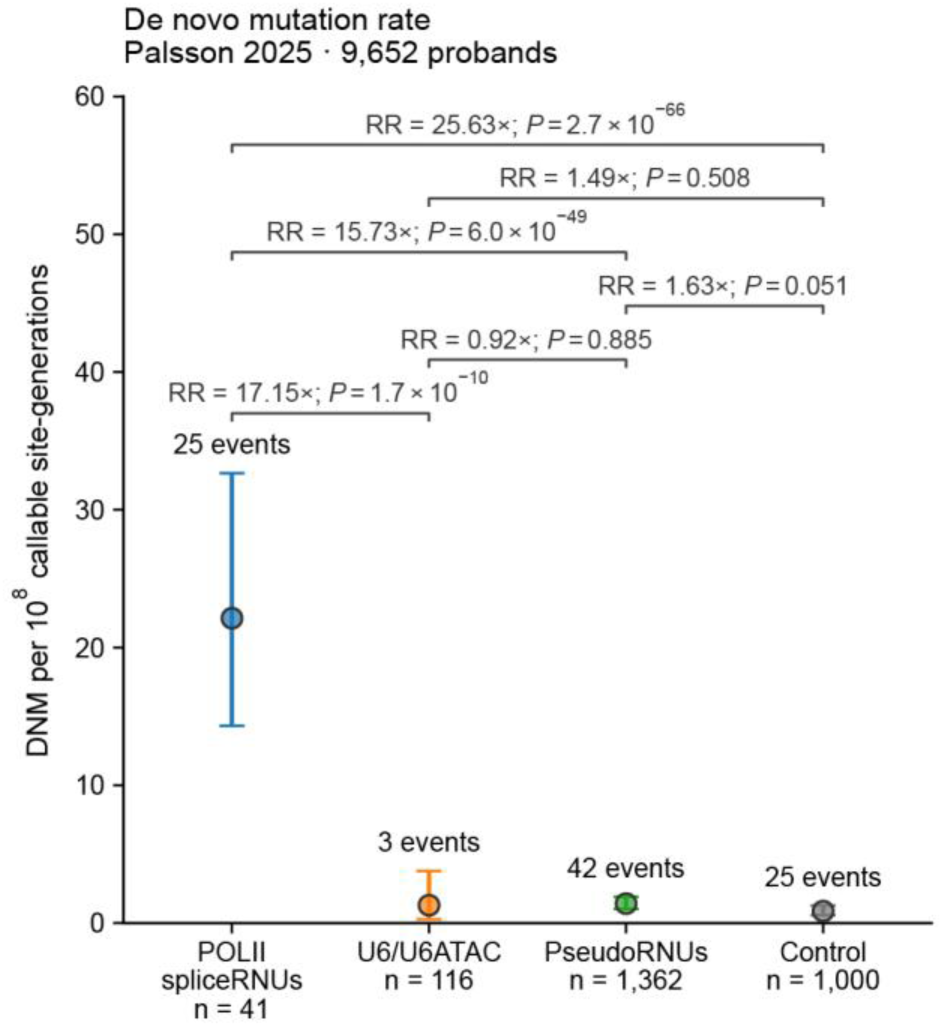
*De novo* mutation rates across different gene groups. *De novo* mutation rates of *Pol II spliceRNUs*, U6/U6ATAC genes, *RNU* pseudogenes and intergenic control regions. Points indicate aggregate rates calculated as the total DNM event count divided by the total exposure, multiplied by 10^8^; exposure is callable base pairs × 9,652 probands × two parental haploid genomes. Error bars denote exact 95% Poisson confidence intervals. The n labels give the numbers of genes or control regions. The brackets report aggregate rate ratios (RR; the rate in the left-hand class divided by that in the right-hand class) and *P* values from two-sided Poisson rate-score tests.

**Supplementary Fig. 4.**
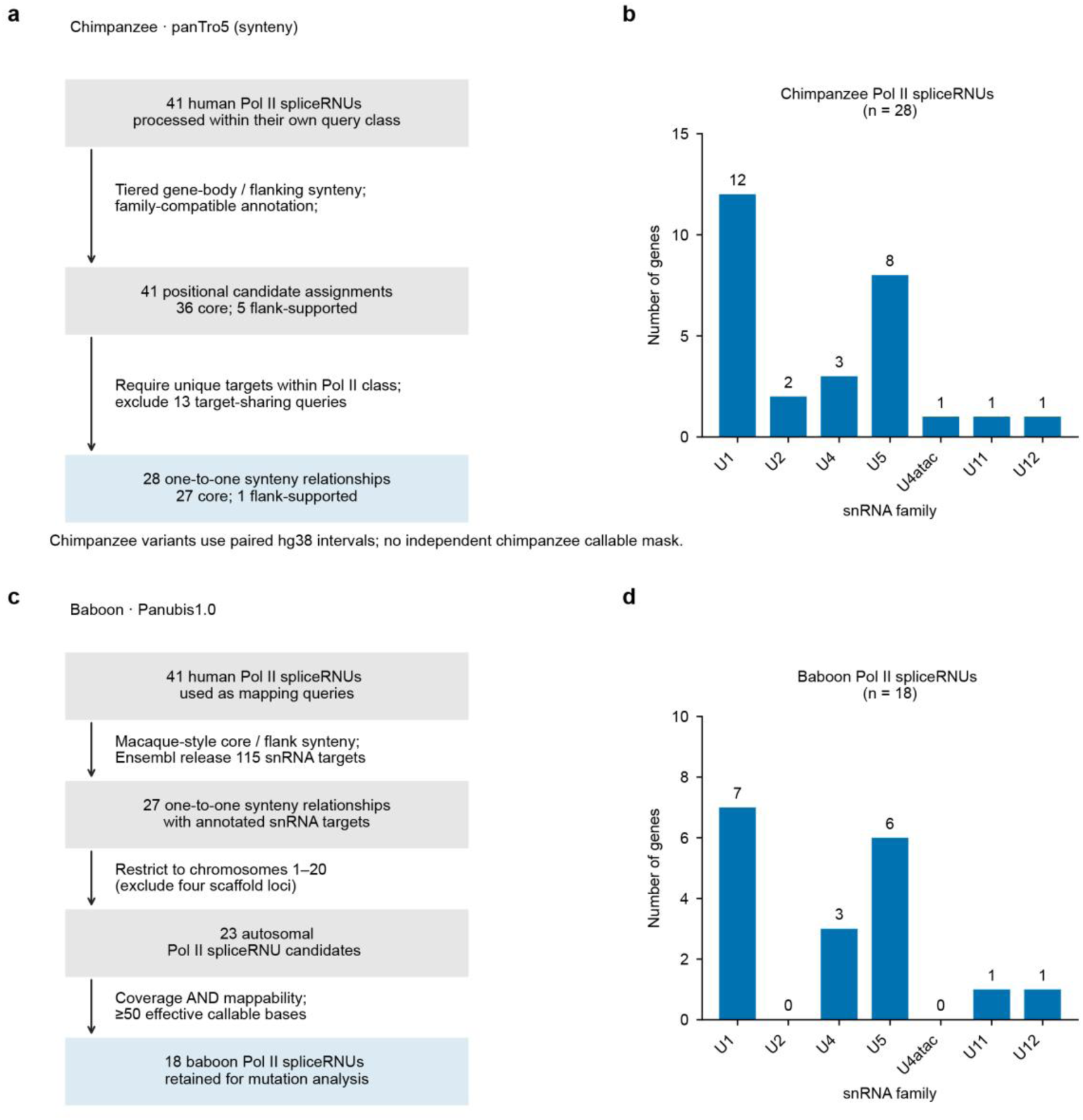
Annotation strategy and family composition of chimpanzee and baboon *Pol II spliceRNUs*. Filtering strategy used to retain genes expressing possible *Pol II spliceRNUs* of chimpanzee (**a**) and baboon (**c**). Number and family composition of chimpanzee (**b**) and baboon (**d**) annotated *Pol II spliceRNUs*, respectively.

**Supplementary Fig. 5.**
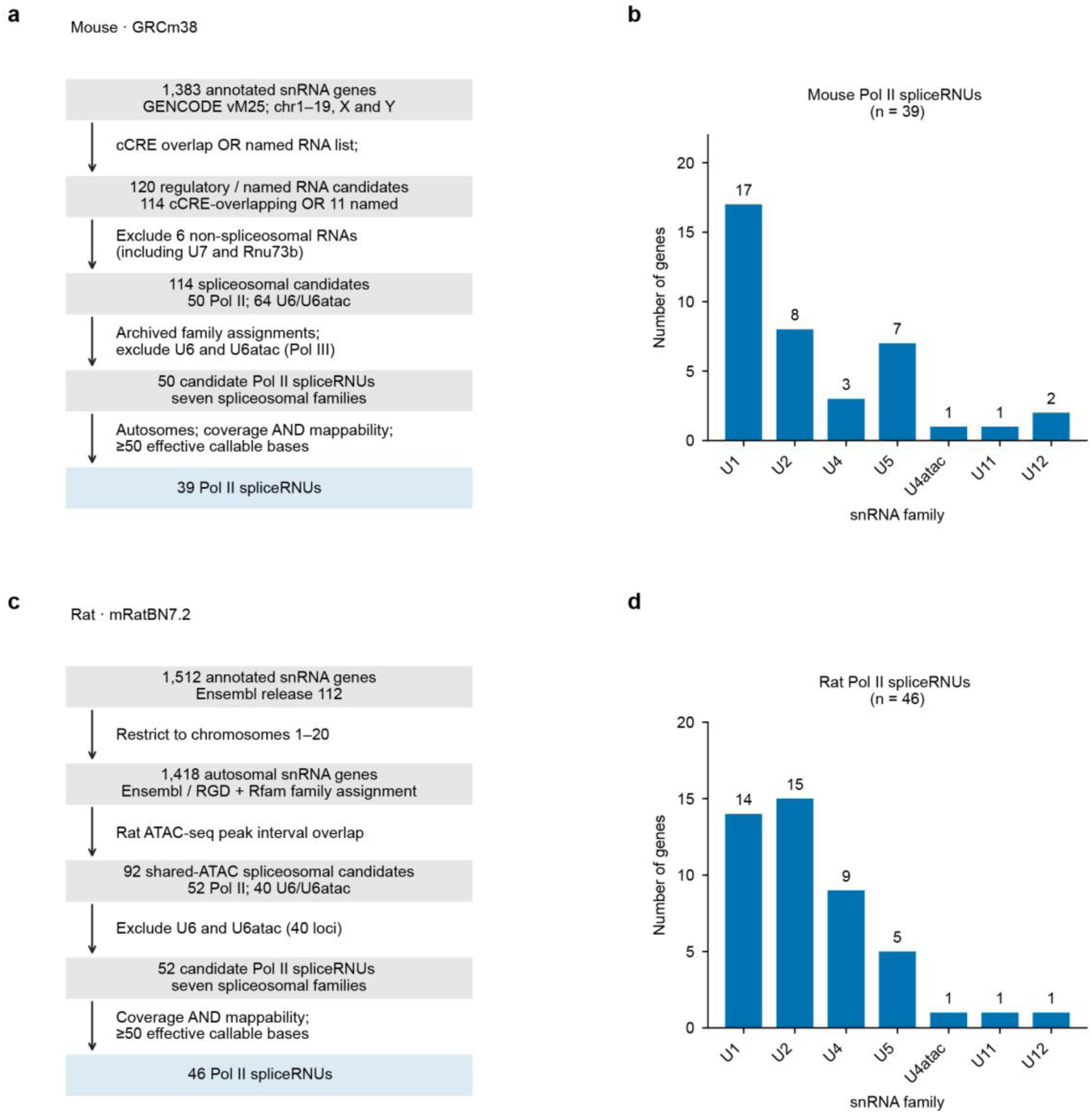
Annotation strategy and family composition of mouse and rat *Pol II spliceRNUs*. Filtering strategy used to retain genes expressing possible *Pol II spliceRNUs* of mouse (**a**) and rat (**c**). Number and family composition of mouse (**b**) and rat (**d**) annotated *Pol II spliceRNUs*, respectively.

**Supplementary Fig. 6.**
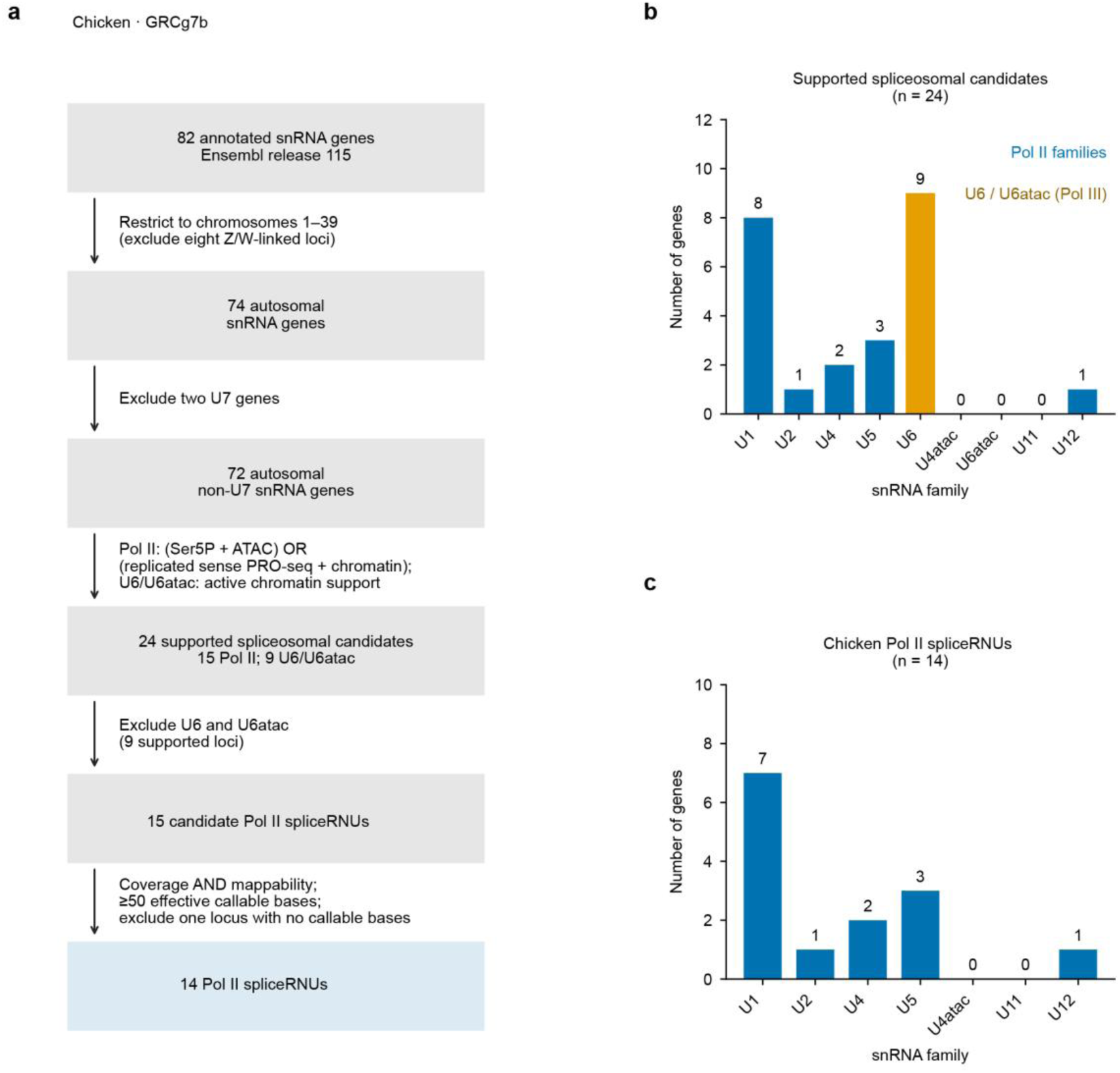
Annotation strategy and family composition of chicken *Pol II spliceRNUs*. **a**, Filtering strategy used to retain genes expressing possible chicken *Pol II spliceRNUs*. **b**, **c**, Number and family composition distribution of chicken annotated *Pol II spliceRNUs* before (**b**) and after (**c**) filtering.

**Supplementary Fig. 7.**
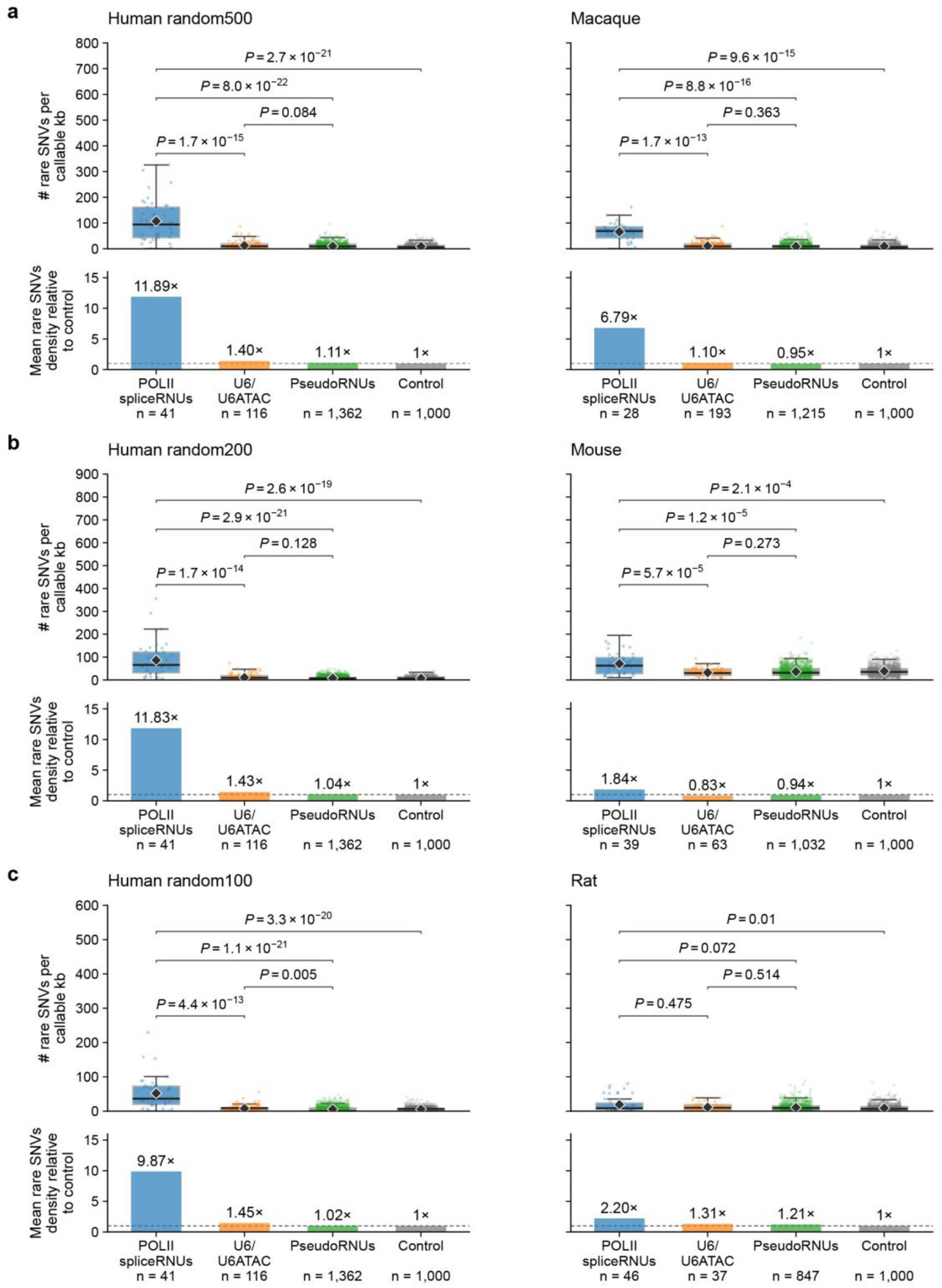
Human downsampling at species-matched allele-frequency thresholds. **a**, Human random500 and macaque (514 individuals), with AF ≤ 2 ×10^-3^. **b**, Human random200 and mouse (148 individuals), with AF ≤ 1 ×10^-2^. **c**, Human random100 and rat (89 individuals), with AF ≤ 2 × 10^-2^. In each panel, the upper row shows gene-level rare SNV density boxplots; the lower row shows the arithmetic-mean density of each class divided by the mean density of control within the same dataset. Human subsets are shown on the left and the corresponding non-human cohorts on the right. These comparisons approximate non-human sample sizes and use the corresponding AF thresholds. Centre lines indicate medians; boxes, interquartile ranges; whiskers, the most extreme observations within 1.5 × the interquartile range; points, individual genes or loci; and diamonds indicate arithmetic means. Brackets compare *Pol II spliceRNUs* with each comparator and report *P* values from two-sided Mann-Whitney *U* tests. Lower bars show mean locus density divided by the corresponding mean control density; dashed lines indicate a ratio of 1. Mean density for each class is divided by the mean density of same-species control loci, defined as 1,000 randomly selected, non-overlapping 150-bp intergenic regions; control equals one by definition.

**Supplementary Fig. 8.**
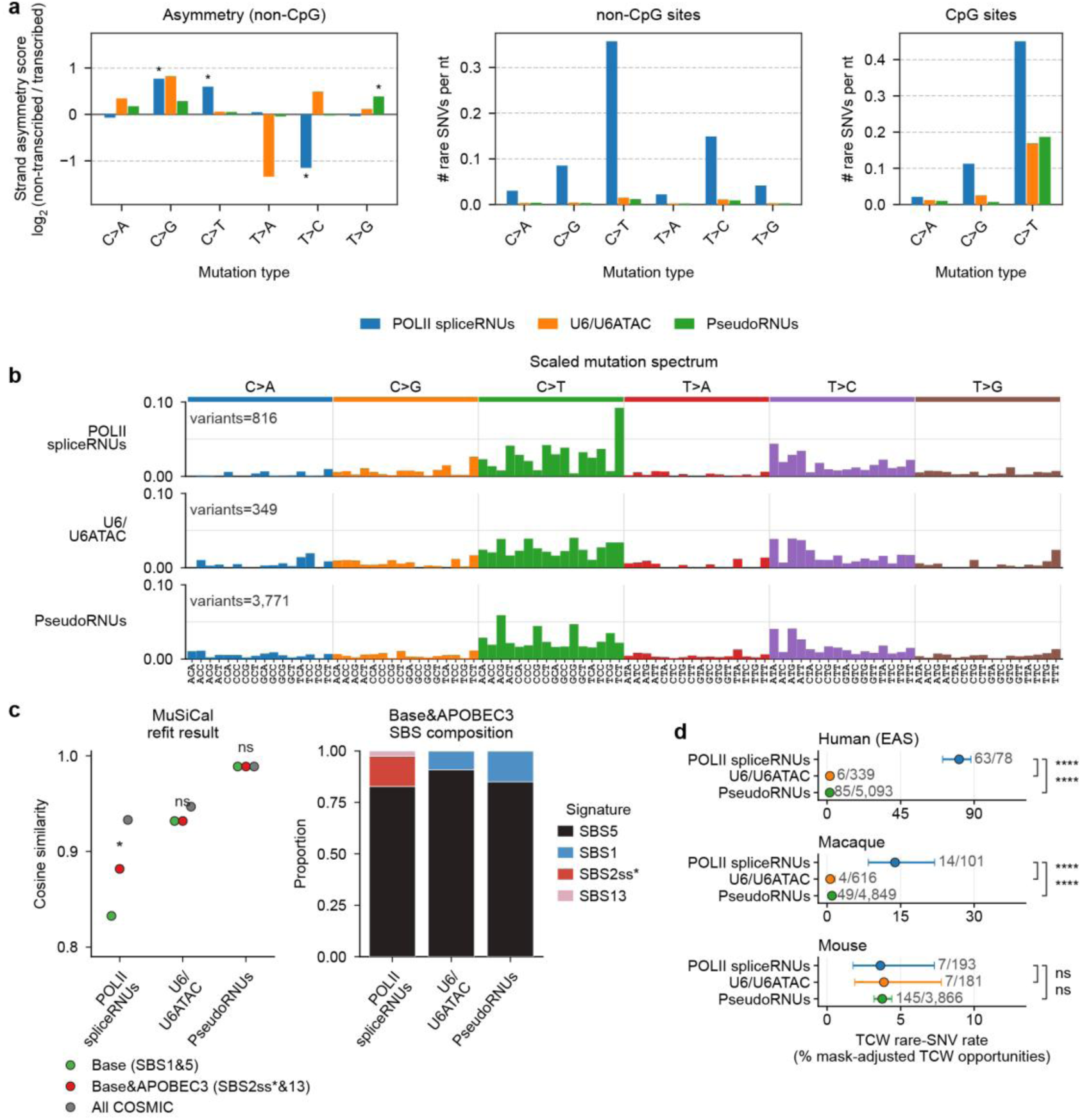
Mutation asymmetry, SBS96 signatures and TCW rare-variant enrichment in the East Asian (EAS) population. **a**, Corrected rare SNV densities (1-mer) and transcriptional asymmetry scores for each gene group in the gnomAD v4 EAS population. CpG and non-CpG sites are shown separately, and asymmetry scores were calculated from non-CpG sites. Asterisks indicate *P* < 0.05 by two-sided Fisher’s exact test. **b**, Scaled mutation spectra for *Pol II spliceRNUs*, U6/U6ATAC genes and *RNU* pseudogenes after normalization to genome-wide 3-mer compositions. The y axis shows the proportion of mutations in each 3-mer context. **c**, MuSiCal refitting under three signature sets (left) and fitted SBS contributions under the Base&APOBEC3 model (right). Asterisks indicate unadjusted empirical *P* < 0.05 from 1,000 targeted permutations; ns, not significant. **d**, Rare SNV rates at TCW sites in *Pol II spliceRNUs*, U6/U6ATAC genes and *RNU* pseudogenes in human EAS, macaque and mouse. Points show observed rates and error bars 95% Wilson confidence intervals. Brackets show *P* values from two-sided Fisher’s exact tests. ns: not significant; *: 0.01 < *P* ≤ 0.05; **: 0.001 < *P* ≤ 0.01; ***: 0.0001 < *P* ≤ 0.001; ****: *P* ≤ 0.0001.

**Supplementary Fig. 9.**
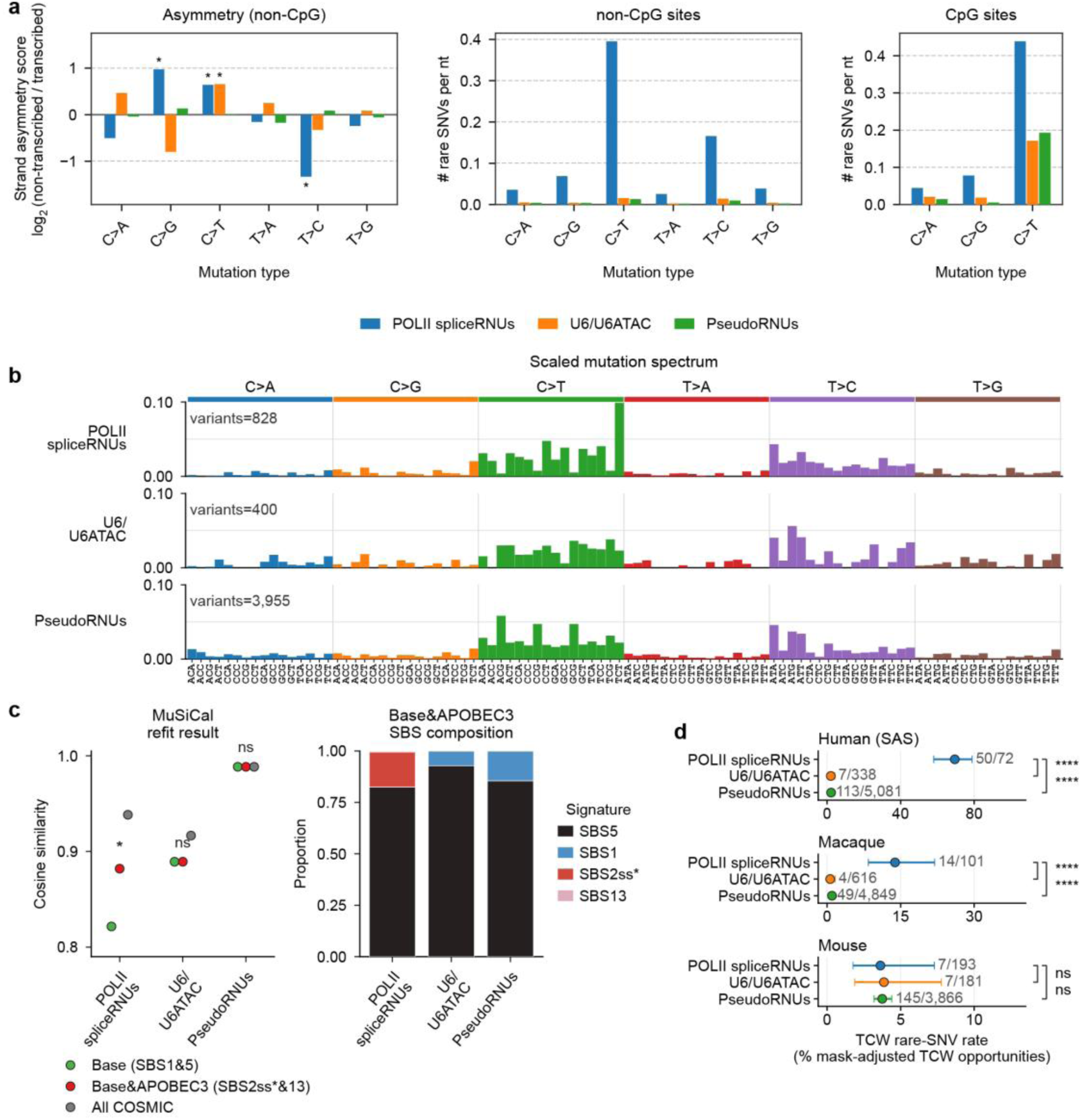
Mutation asymmetry, SBS96 signatures and TCW rare-variant enrichment in the South Asian (SAS) population. **a**, Corrected rare SNV densities (1-mer) and transcriptional asymmetry scores for each gene group in the gnomAD v4 SAS population. CpG and non-CpG sites are shown separately, and asymmetry scores were calculated from non-CpG sites. Asterisks indicate *P* < 0.05 by two-sided Fisher’s exact test. **b**, Scaled mutation spectra for *Pol II spliceRNUs*, U6/U6ATAC genes and *RNU* pseudogenes after normalization to genome-wide 3-mer compositions. The y axis shows the proportion of mutations in each 3-mer context. **c**, MuSiCal refitting under three signature sets (left) and fitted SBS contributions under the Base&APOBEC3 model (right). Asterisks indicate unadjusted empirical *P* < 0.05 from 1,000 targeted permutations; ns, not significant. **d**, Rare SNV rates at TCW sites in *Pol II spliceRNUs*, U6/U6ATAC genes and *RNU* pseudogenes in human SAS, macaque and mouse. Points show observed rates and error bars 95% Wilson confidence intervals. Brackets show *P* values from two-sided Fisher’s exact tests. ns: not significant; *: 0.01 < *P* ≤ 0.05; **: 0.001 < *P* ≤ 0.01; ***: 0.0001 < *P* ≤ 0.001; ****: *P* ≤ 0.0001.

**Supplementary Fig. 10.**
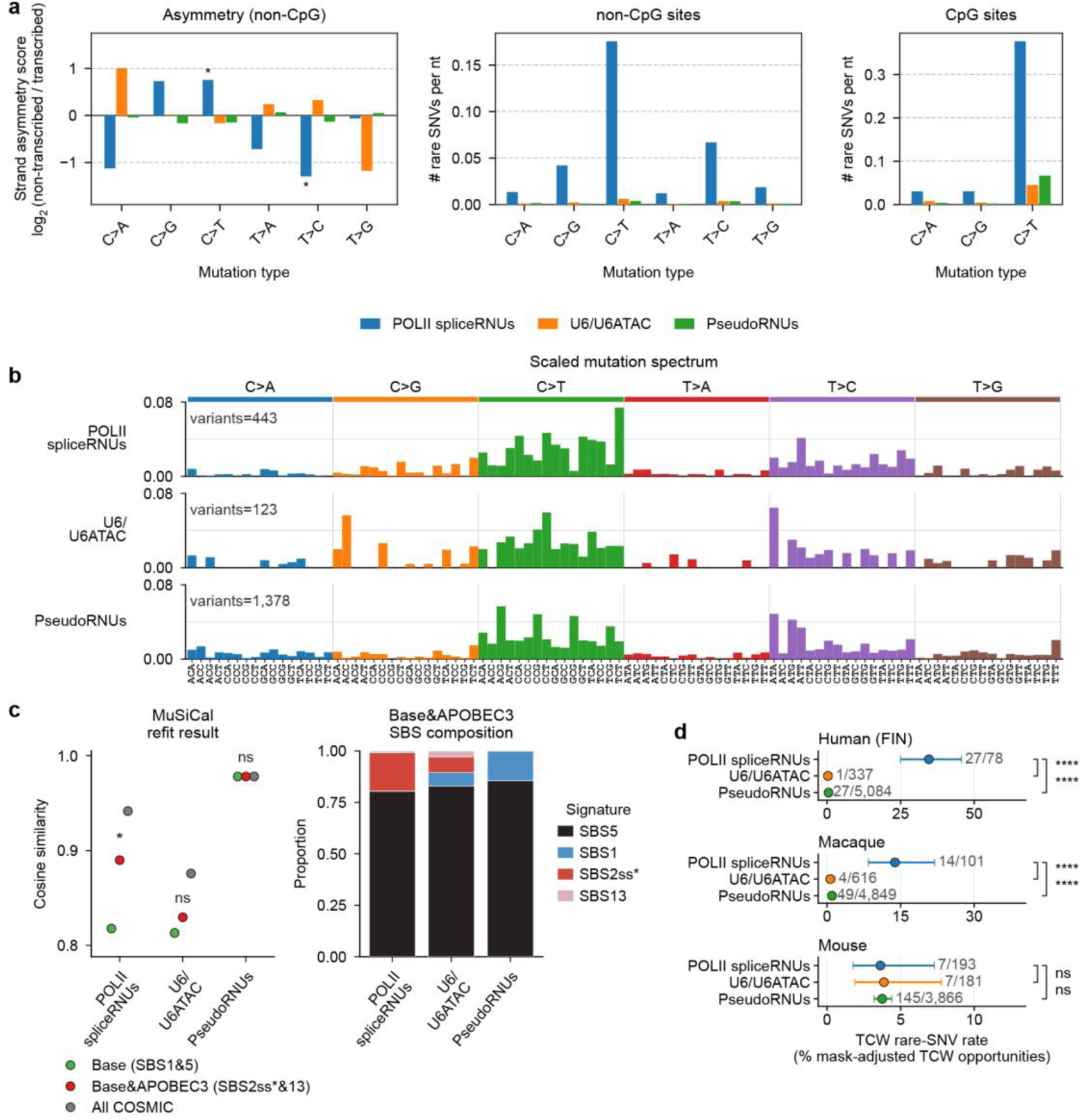
Mutation asymmetry, SBS96 signatures and TCW rare-variant enrichment in the Finnish (FIN) population. **a**, Corrected rare SNV densities (1-mer) and transcriptional asymmetry scores for each gene group in the gnomAD v4 FIN population. CpG and non-CpG sites are shown separately, and asymmetry scores were calculated from non-CpG sites. Asterisks indicate *P* < 0.05 by two-sided Fisher’s exact test. **b**, Scaled mutation spectra for *Pol II spliceRNUs*, U6/U6ATAC genes and *RNU* pseudogenes after normalization to genome-wide 3-mer compositions. The y axis shows the proportion of mutations in each 3-mer context. **c**, MuSiCal refitting under three signature sets (left) and fitted SBS contributions under the Base&APOBEC3 model (right). Asterisks indicate unadjusted empirical *P* < 0.05 from 1,000 targeted permutations; ns, not significant. **d**, Rare SNV rates at TCW sites in *Pol II spliceRNUs*, U6/U6ATAC genes and *RNU* pseudogenes in human FIN, macaque and mouse. Points show observed rates and error bars 95% Wilson confidence intervals. Brackets show *P* values from two-sided Fisher’s exact tests. ns: not significant; *: 0.01 < *P* ≤ 0.05; **: 0.001 < *P* ≤ 0.01; ***: 0.0001 < *P* ≤ 0.001; ****: *P* ≤ 0.0001.

**Supplementary Fig. 11.**
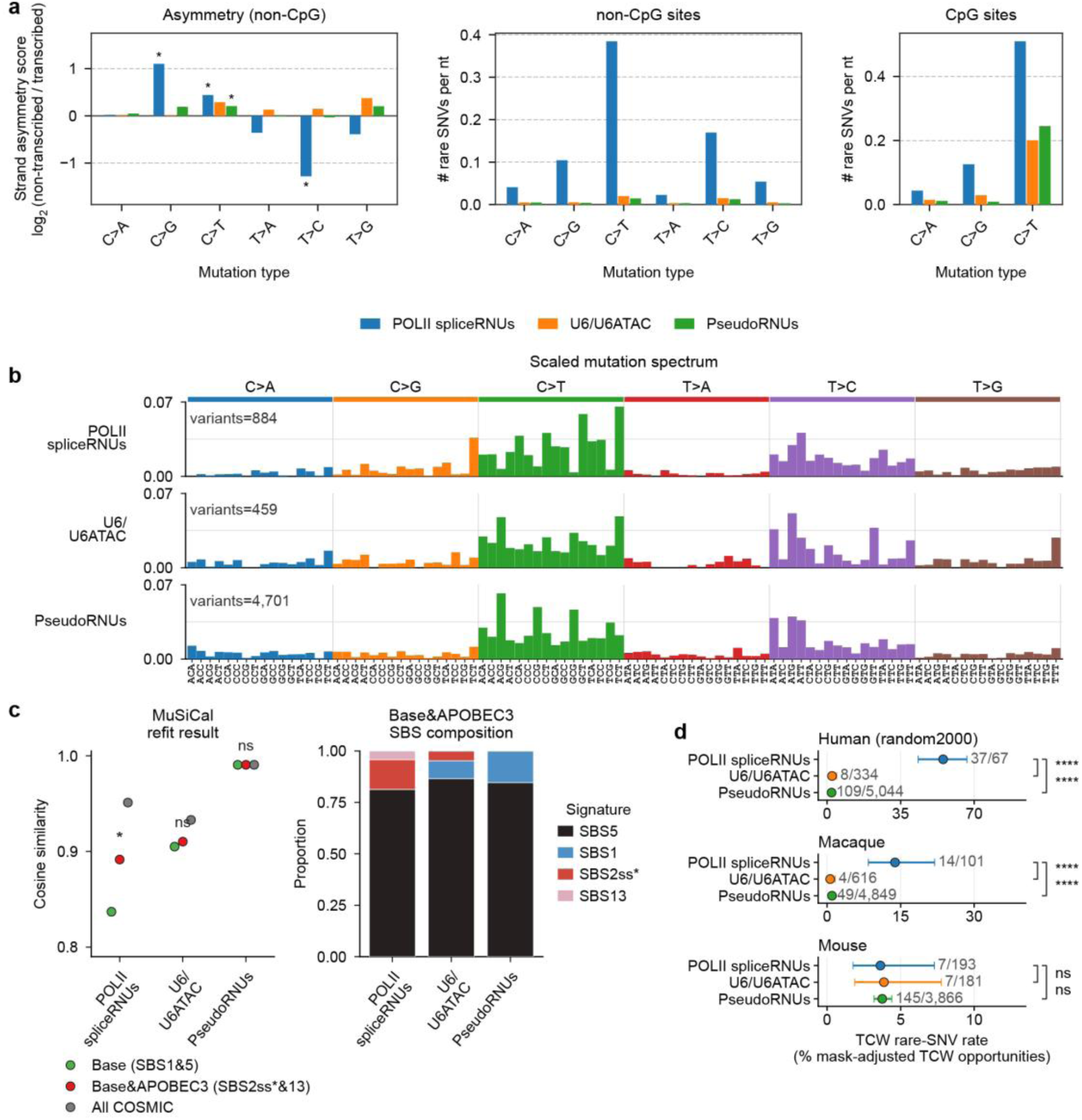
Mutation asymmetry, SBS96 signatures and TCW rare-variant enrichment in the randomly sampled 2,000-individual (random2000) cohort. **a**, Corrected rare SNV densities (1-mer) and transcriptional asymmetry scores for each gene group in the random 2,000-individual HGDP+1KG subset (random2000) population. CpG and non-CpG sites are shown separately, and asymmetry scores were calculated from non-CpG sites. Asterisks indicate *P* < 0.05 by two-sided Fisher’s exact test. **b**, Scaled mutation spectra for *Pol II spliceRNUs*, U6/U6ATAC genes and *RNU* pseudogenes after normalization to genome-wide 3-mer compositions. The y axis shows the proportion of mutations in each 3-mer context. **c**, MuSiCal refitting under three signature sets (left) and fitted SBS contributions under the Base&APOBEC3 model (right). Asterisks indicate unadjusted empirical *P* < 0.05 from 1,000 targeted permutations; ns, not significant. **d**, Rare SNV rates at TCW sites in *Pol II spliceRNUs*, U6/U6ATAC genes and *RNU* pseudogenes in human random2000, macaque and mouse. Points show observed rates and error bars 95% Wilson confidence intervals. Brackets show *P* values from two-sided Fisher’s exact tests. ns: not significant; *: 0.01 < *P* ≤ 0.05; **: 0.001 < *P* ≤ 0.01; ***: 0.0001 < *P* ≤ 0.001; ****: *P* ≤ 0.0001.

## Supplementary Tables

Titles of Supplementary Tables are shown below. See details in the associated Excel file.

Supplementary Table 1. Datasets used in snRNA annotation of representative species.

Supplementary Table 2. Annotations of human spliceosomal snRNA genes and pseudogenes.

Supplementary Table 3. Annotations of chimpanzee spliceosomal snRNA genes and pseudogenes and syntenic locations in human GRCh38.

Supplementary Table 4. Annotations of macaque spliceosomal snRNA genes and pseudogenes.

Supplementary Table 5. Annotations of baboon spliceosomal snRNA genes and pseudogeness.

Supplementary Table 6. Annotations of mouse spliceosomal snRNA genes and pseudogenes.

Supplementary Table 7. Annotations of rat spliceosomal snRNA genes and pseudogenes.

Supplementary Table 8. Annotations of chicken spliceosomal snRNA genes and pseudogenes.

Supplementary Table 9. Source of APOBEC3 repertoire annotations.

Supplementary Table 10. Corrected rare SNV densities (1-mer) for in Pol II spliceRNUs, U6/U6ATAC genes and RNU pseudogenes in human.

Supplementary Table 11. Calculation of transcriptional asymmetry scores for each gene group in the gnomAD v4 populations.

Supplementary Table 12. Mutation signature analysis under different variant filtering and processing strategies.

Supplementary Table 13. Rare SNV rates at TCW sites in Pol II spliceRNUs, U6/U6ATAC genes and RNU pseudogenes in human populations, macaque and mouse.

Supplementary Table 14. Functional genomics datasets used for analysis in Fig. 3.

Supplementary Table 15. Fractions of genes overlapping human RLBase regions or the union of nine public mouse embryonic stem-cell R-loop peak sets.

Supplementary Table 16. List of species, genome assemblies, and population datasets used for analysis of snRNA genes.

